# A conserved nuclear export complex coordinates transcripts for dopaminergic synaptogenesis and neuronal surviva

**DOI:** 10.1101/282137

**Authors:** Celine I. Maeder, Jae-Ick Kim, Konstantin Kaganovsky, Ao Shen, Qin Li, Zhaoyu Li, X.Z. Shawn Xu, Jin Billy Li, Yang K. Xiang, Jun B. Ding, Kang Shen

## Abstract

Synaptic vesicle and active zone proteins are required for synaptogenesis. The molecular mechanisms for coordinated synthesis of these proteins are not understood. Using forward genetic screens, we identified the conserved <u>THO</u> nuclear export <u>C</u>omplex (THOC) as master regulator of presynapse development in *C.elegans* dopaminergic neurons. In THOC mutants, synaptic messenger RNAs are trapped in the nucleus, resulting in dramatic decrease of synaptic protein expression, near complete loss of synapses and compromised dopamine function. cAMP-responsive element binding protein (CREB) interacts with THOC to mark activity-dependent transcripts for efficient nuclear export. Deletion of the THOC subunit Thoc5 in mouse dopaminergic neurons causes severe defects in synapse maintenance and subsequent neuronal death in the Substantia Nigra compacta (SNc). These cellular defects lead to abrogated dopamine release, ataxia and animal death. Together, our results argue that nuclear export mechanisms can select specific mRNAs and be a rate-limiting step for synapse development and neuronal survival.

**Highlights:**

- Dopaminergic presynapses are severely impaired in *thoc* mutant worms and mice
- THOC specifically controls the nuclear export of synaptic transcripts
- CREB recruits THOC onto activity-dependent synaptic transcripts for efficient export
- Dopamine neurons in the SNc degenerate upon conditional knock-out of *thoc5*

## Introduction

Neurons communicate with each other through the synapse, a highly specialized structure for signal transmission and reception. Synapses are assembled during neuron development. They are built from hundreds of proteins (Laßek et al., 2015), which assemble into functionally distinct pre- and postsynaptic compartments. While the molecular composition of pre- and postsynapses has been widely explored through proteomic studies (Laßek et al., 2015), and significant progress has been achieved in identifying the key molecular players in the physical process of synapse assembly (Frank and Grant, 2017; Patel et al., 2006; Van Vactor and Sigrist, 2017), much less is known, how synaptogenesis is regulated at the level of gene expression. Is there a mechanism that coordinates the expression of functionally related proteins (e.g. synaptic proteins) such that they are ready to assemble into higher order structures concomitantly?

Regulation of gene expression has been shown to be important for neuronal development and function under many circumstances. Gene expression is a multi-step process starting with the transcription of messenger RNA (mRNA), followed by mRNA maturation, nuclear export and translation in the cytoplasm. Regulation of gene expression happens at every step along the process. At the level of gene transcription, terminal selector transcription factors initiate and maintain the terminal differentiation program of neuronal subtypes by controlling the expression of neuron-type specific gene batteries (Hobert, 2016). Similarly, the cAMP-responsive element binding protein (CREB) transcription factor couples neuronal activity to long-term changes in synaptic plasticity and memory formation by inducing the transcription of immediate early genes (IEGs) (Sakamoto et al., 2011). Interestingly, multiple studies found that pan-neuronal genes are not regulated by a single transcription factor; they rather contain redundant parallel-acting cis-regulatory modules in their promoters, which are responsive to multiple trans-acting factors (Liu et al., 2009; Stefanakis et al., 2015). Splicing adds another layer of gene expression regulation. For example, alternative splicing of cell adhesion molecules such as the mammalian neurexins and the *Drosophila* Dscam was shown to regulate neuronal cell recognition and the establishment of specific neuronal circuits (Raj and Blencowe, 2015). At the level of protein translation, local translation has emerged as an elegant way to spatially regulate protein synthesis within a neuron. A large body of literature demonstrates that in dendrites, synaptic activity induces local translation of so-called plasticity related genes to facilitate synaptic plasticity and memory formation (Rangaraju et al., 2017) and that perturbations in this process may contribute to neurodevelopmental and neurodegenerative disease (Iacoangeli and Tiedge, 2013; Lenzken et al., 2014). Local translation has also been reported in developing axons, where it contributes to regulation of axon path finding (Rangaraju et al., 2017). Lastly, gene expression can be modulated at the level of nuclear export of mRNAs to the cytoplasm. Importantly, multiple studies demonstrated that defects in nuclear mRNA export might be an underlying cause of neurodegenerative disease such as in G4C2 repeat expansion of C9orf72 amyotrophic lateral sclerosis (ALS) or polyglutamine disease (Freibaum et al., 2015; Tsoi et al., 2011; Zhang et al., 2015). Furthermore, mutations in the mRNA export mediator Gle1 cause fetal motoneuron disease and gle1 mutant zebrafish embryos show dramatic mRNA accumulation in the nucleus (Nousiainen et al., 2008; Seytanoglu et al., 2016). Less is known which RNA binding proteins (RBPs) are involved in the regulation of mRNA export in neurons and whether different RBPs might regulate the export of unique subsets of functionally related mRNAs. In non-neuronal cells, Botti et al. demonstrated that the cellular differentiation state can modulate the mRNA export activity of certain SR proteins through distinct posttranslational modifications on these SR proteins (Botti et al., 2017).

The THO complex (THOC) is a conserved RNA binding complex which has been implicated in export of subsets of mRNAs in diverse biological processes (Guria et al., 2011; Saran et al., 2013; Tran et al., 2013, 2014a; Wang et al., 2013). Together with the RNA helicase UAP56 and the export adaptor AlyRef/THOC4, it forms the TREX (TRanscription/EXport) complex (Sträßer et al., 2002) that facilitates the formation of export-competent messenger ribonucleoprotein complexes (mRNPs) by recruiting additional mRNA processing and exporting factors (Heath et al., 2016). Even though THOC’s core function in mRNA maturation and export seems to be conserved from yeast to mammals, it has diverse cellular functions in different cell types. In *S. cerevisae*, the THOC consists of Hrp1, Tho2, Mft1, Thp2 and Tex1 and functions in RNA transcription, nuclear export and genome stability (Jimeno et al., 2002; Sträßer et al., 2002). In higher eukaryotes such as *Drosophila* (Rehwinkel et al., 2004) and humans (Masuda et al., 2005), the THOC is composed of THOC1, THOC2 and THOC3, the human homologs of Hpr1, Tho2 and Tex1, respectively, as well as three additional unique proteins THOC5, THOC6 and THOC7. The THOC is involved in embryonic, hematopoietic and intestinal stem cell renewal and survival, testis development as well as cell proliferation in cancer (Mancini et al., 2010; Pitzonka et al., 2014; Saran et al., 2016, 2013; Tran et al., 2013; Wang et al., 2013, 2009). Interestingly, THOC appears to regulate different mRNA species in diverse cell types suggesting that its function is tailored to achieve context-or cell type-specific gene expression programs. Only little is known about THOC’s function in neurodevelopment and neurodegenerative disease. Missense mutations in the *THOC2* gene were shown to cause syndromic intellectual disability (Kumar et al., 2015). Additionally, a de-novo translocation close to the *THOC2* gene was reported in a child with cerebellar hypoplasia, ataxia and retardation. Impaired movement was also seen in *thoc-2* mutant *C. elegans* (Di Gregorio et al., 2013). In case of *THOC6*, multiple bi-allelic mis-/non-sense mutations have been associated with intellectual disability, brain malformation as well as renal and heart defects (Amos et al., 2017; Anazi et al., 2016; Beaulieu et al., 2013; Boycott et al., 2010). Some of these mutations caused mislocalization of THOC6 from the nucleus to the cytoplasm, suggesting an important nuclear role for THOC during neurodevelopment. It is noteworthy to mention that the THOC is highly enriched and mis-localized to cytoplasmic protein aggregates, a hallmark of neurodegenerative disease, in primary neurons (Woerner et al., 2016). It also colocalizes with cytoplasmic huntingtin protein aggregates in the R6/2 huntingtin disease mouse model. These data suggest that impairment in nucleocytoplasmic transport in part facilitated by THOC may contribute to the cellular pathology of neurodegenerative disease.

In this study we identified the THOC as a master regulator of presynaptic gene expression in *C. elegans* through an unbiased forward genetic screen. THOC mutants have strong defects in dopaminergic synaptogenesis resulting in near complete loss of synapses, impaired dopamine (DA) signaling and corresponding behavioral deficits. Using single molecule fluorescence *in situ* hybridization (smFISH), we show that presynaptic transcripts, but not control transcripts, are trapped in the nucleus of DA neurons, leading to a depletion of their mRNAs in the cytoplasm and hence reduced protein synthesis. Furthermore, we demonstrate that THOC’s function in presynaptic gene expression is especially pivotal under neuronal activity. Based on our genetic as well as biochemical data we propose a model, whereby the activity-dependent transcription factor CREB recruits THOC to presynaptic transcripts, hence facilitating their efficient export out of the nucleus, allowing for concerted translation, which ultimately enables a temporally coordinated assembly of dopaminergic presynapses.

Lastly, using conditional *thoc5* knock-out mice, we demonstrate that THOC’s function in neurons is conserved from worms to mammals. Mice homozygous mutant for *thoc5* just in DA neurons develop normally up to 3 weeks of life, when they start to lose dopaminergic synapses. At 6 weeks of age, most DA neurons die, especially in the substantia nigra pars compacta (SNc), leading to sever impairment of DA release and locomotion defects in these animals. Taken together, our data demonstrate that the evolutionarily conserved THOC plays an essential role in synaptogenesis by regulating the export of synaptic transcripts.

## Results

### Dopaminergic presynaptic specializations are severely impaired in THOC mutant animals

*C. elegans* PDEL and PDER are a pair of dopaminergic neurons with their cell bodies in the posterior part of the animal. They both extend sensory, ciliated dendrites into the cuticle and project their axons ventrally, where they bifurcate and run along the entire ventral nerve cord (Figure 1A and (White et al., 1986)). They form evenly spaced *en-passant* synapses along the axon, which can be visualized by the expression of fluorescently tagged presynaptic components under a DA neuron specific promoter (Figure 1A, B).

**Figure 1:**
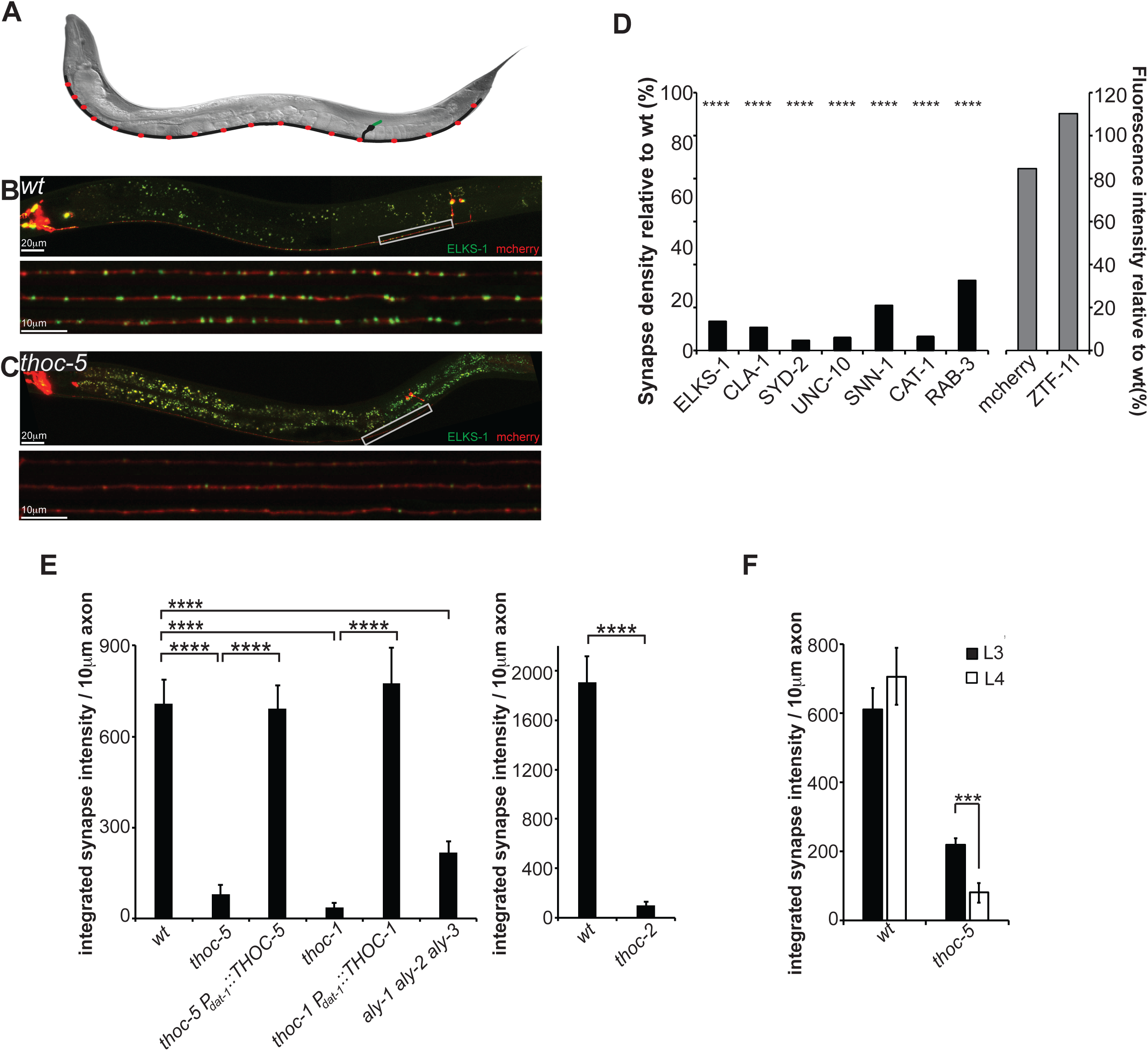
Dopaminergic presynapses are severely impaired in *thoc* mutant animals. (A) Schematic of the dopaminergic PDE neuron in *C. elegans* (green: dendrite; black: cellbody and axon; red: presynapses) (B) Image of wildtype PDE neuron *in vivo* at larval stage L4, visualized by the expression of cytosolic mCherry and GFP::ELKS-1 under a DA-neuron specific promoter. Lower panel: Line scans of three different PDE axons. (C) Representative image of a *thoc-5* mutant animal at larval stage L4. Lower panel: Line scans of three different PDE axons showing dramatic decrease in presynapses. (D) Multiple presynaptic markers (black bars) are significantly downregulated in *thoc-5* mutant animals, however control proteins (grey bars) are unaffected. N=5-20 animals at L4 stage; ^****^, p<0.0001, one-way ANOVA with Tukey post-hoc test. (E) Quantification of *thoc-1, thoc-2, thoc-5* single mutant and *aly-1 aly-2 aly-3* triple mutant phenotypes. Cell-specific expression of THOC-1 and THOC-5 rescues the impairment of DA presynapses. N=10-20 animals at L4 stage; ^****^, p<0.0001, one-way ANOVA with Tukey post-hoc test. (F) Synapse defect aggravates with age. Dopaminergic presynapses were quantified at larval stages L3 and L4 in *wildtype* and *thoc-5* mutant animals. N=10-20 animals; ^***^, p<0.001, student t-test. Averages and SEM are plotted in (E) and (F). See also Figure S1.

To identify molecules important for the concerted expression of presynaptic proteins, we performed a visual forward genetic screen and isolated two mutations *wy817* and *wy822*, with dramatic defects in presynaptic specializations, however intact axonal morphology (as assayed by the active zone protein GFP::ELKS-1 and cytoplasmic mCherry, Figure 1B). In both mutants, GFP::ELKS-1 was dramatically reduced and the animals retained only 10 percent of *wildtype* synapses, with the remaining synapses being smaller and dimmer (Figure 1C, D, E and Figure S1). To understand whether these mutations affected the expression of other presynaptic components, we crossed *wy822* to an array of marker strains expressing fluorescently tagged active zone molecules (CLA-1/piccolo; SYD-2/liprinα, UNC-10/RIM), synaptic vesicle proteins (SNN-1/synapsin; CAT-1/VMAT; RAB-3/RAB3) or control proteins. While all presynaptic markers showed a dramatic decrease in synaptic density and size in the mutant background (Figure 1D), control proteins such as cytosolic mCherry or the pro-neurogenic transcription factor ZTF-11/MYT1L did not show any difference in fluorescence intensity between *wildtype* and the mutant (Figure 1D).

Using single nucleotide polymorphism mapping and whole genome sequencing, we identified *wy817* as a splice acceptor site mutation in the 11^th^ intron of *THOC-1*, while *wy822* introduced an early stop codon in *THOC-5* (R288stop). THOC-1 and THOC-5 are part of a nuclear, multi-protein complex conserved from yeast to mammals, which has been implicated to work at the interface of transcription and mRNA export (Chávez et al., 2000; Sträßer et al., 2002).

Strains carrying either a deletion in *THOC-*2 or triple mutations in all *C. elegans* orthologues of THOC4/AlyRef (*aly-1 aly-2 aly-3*) showed similar presynaptic defects as *thoc-1* and *thoc-5* mutants (Figure 1E). While *thoc-5* single and *aly-1 aly-2 aly-3* triple mutant animals are viable as homozygous mutants, *thoc-1* and *thoc-2* mutants are sterile and lethal at young adult stage, hence can only be maintained as heterozygous animals. Using cell-specific expression of THOC-1 or THOC-5 cDNA in the corresponding mutant strain background, we show that THOC functions cell-autonomously in DA neurons (Figure 1E). However, DA neuron specific expression of THOC-1 does not rescue the lethality phenotype, suggesting that this complex also functions in other tissues.

To address whether THOC functions in initial DA synapse development or synapse maintenance, we quantified dopaminergic presynapses at different larval stages. Interestingly, we found that *thoc-5* mutant worms at larval L3 stage contained more abundant and brighter synapses as compared to larval L4 stage (Figure 1F and Figure S1). In contrast, synapse density did not change with age under *wildtype* conditions and synapse intensity showed a slight trend of growth and maturation at L4 as compared to L3 (Figure 1F and Figure S1).

Taken together, our forward genetic screen identified the THOC as an important regulator of presynaptic gene expression in DA neurons.

### THOC also affects other types of presynapses, however to a lesser extent

Next we wanted to explore whether the THOC plays a role in presynaptic gene expression of all neurons or whether it only affects DA neurons. First, we assayed endogenous presynaptic protein levels by quantitative western blot. We examined five presynaptic proteins including SNB-1/synaptobrevin, SNG-1/synaptogarin, SNT-1/synaptotagmin, UNC-64/syntaxin and ELKS-1/ELKS. The antibodies were specific, as we did not detect any bands in corresponding mutant strains (Δ lane). Interestingly, all of the proteins except SNT-1 showed a mild reduction in the *thoc-5* mutants compared to controls (Figure 2A). For ELKS-1, we examined the expression level in a transgenic strain expressing GFP::ELKS-1 in the DA neurons, so that both the endogenous pan-neuronal ELKS-1 as well as the dopamine-specific GFP::ELKS-1 could be assayed simultaneously. Endogenous ELKS-1 was affected to a much milder extent as compared to dopaminergic GFP::ELKS-1 (Figure 2A).

**Figure 2:**
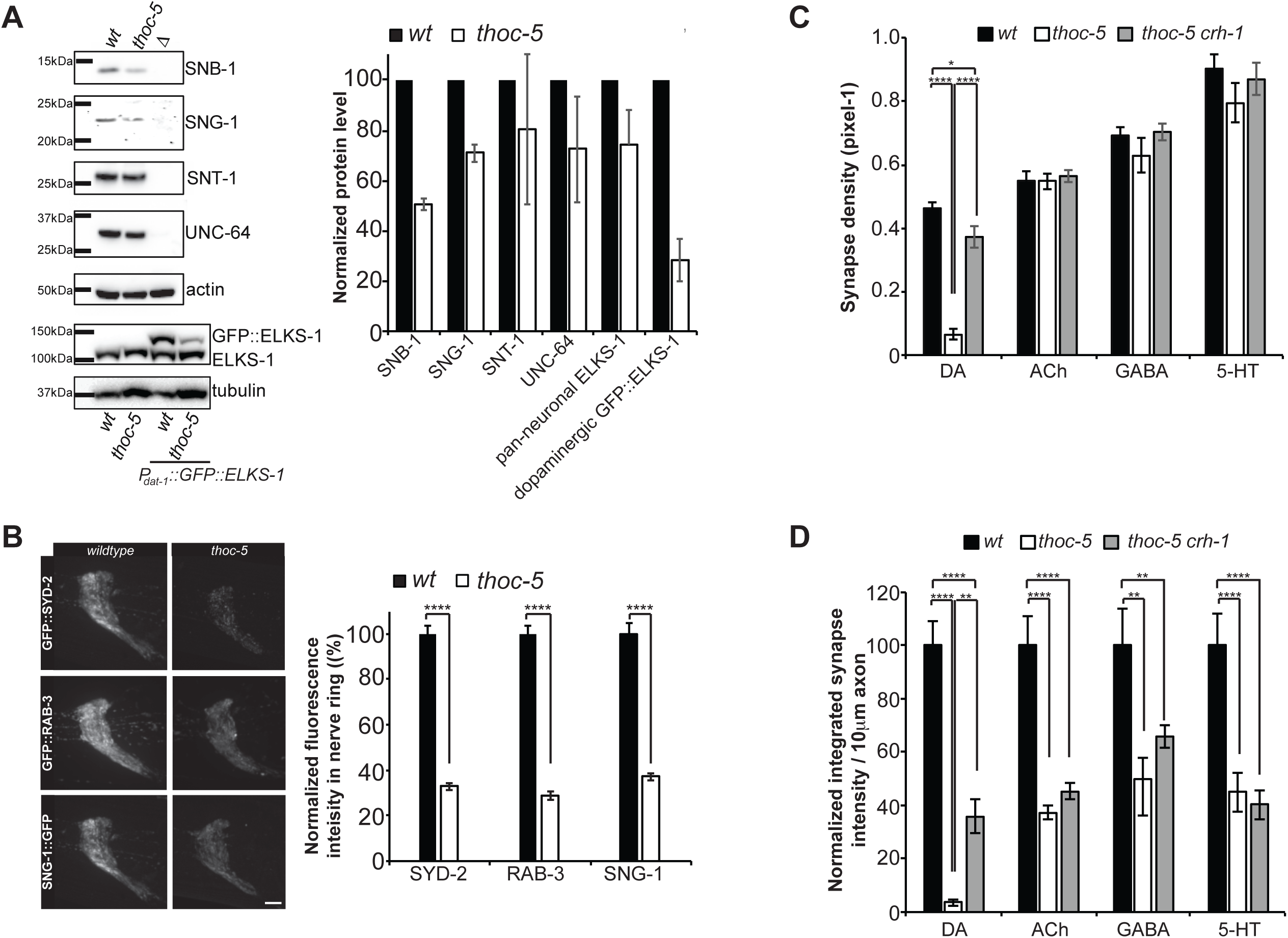
THOC most severely affects dopamine synapses. (A) Quantitative western blotting of endogenous, pan-neuronally expressed presynaptic proteins. Representative blot images and quantification of three independent experiments is shown. (B) Endogenous GFP knock-in strains for SYD-2, RAB-3 and SNG-1. Representative images of all the synapses in the nerve ring is shown for *wildtype* and *thoc-5* mutant animals. Quantification of total nerve ring fluorescence is plotted. N=10-20 animals. ^****^, p<0.0001, student t-test. Scale bar represents 5□m. (C) Synapse density quantification for different types of neurons in *wildtype, thoc-5* and *thoc-5 crh-1* mutant backgrounds. N=10-20 animals at L4 stage; ^****^, p<0.0001, one-way ANOVA with Tukey post-hoc test. (D) Normalized integrated synaptic intensity is quantified for different types of neurons in *wildtype, thoc-5* and *thoc-5 crh-1* mutant backgrounds. N=10-20 animals at L4 stage; ^****^, p<0.0001; ^**^, p<0.01, one-way ANOVA with Tukey post-hoc test. Averages and SEM are plotted.

To further test if synaptically localized SV and AZ proteins are affected by *thoc-5 in vivo*, we generated knock-in strains, where GFP was inserted into the endogenous genetic loci of *syd-2*/liprinα, *rab-3*/RAB3 and *sng-1*/synaptogyrin with CRISPR/Cas9. These knock-in strains showed punctate GFP fluorescent signals in the neuropils. The strongest signals were found in the “nerve ring” structure, which is the biggest neuropil in *C. elegans* with more than 100 axons (8 of which are dopaminergic). Since the nerve ring does not contain any neuronal cell bodies, GFP fluorescence at the nerve ring reflects synaptically localized proteins. We quantified the total fluorescence intensity of these proteins in the nerve ring in *wildtype* and *thoc-5* mutant background. Again, we found a global decrease in fluorescence intensity in *thoc* mutant animals for all three synaptic proteins (Figure 2B).

Lastly, to directly study THOC’s function in different neuronal types, we utilized a cell-type specific endogenous GFP knock-in strain (Schwartz and Jorgensen, 2016), in which the tagging of the SV protein RAB-3 with GFP at the endogenous locus was activated by the expression of flippase under different neuron-type specific promoters. Using this labeling strategy, we found that cholinergic, GABAergic and serotonergic synapses did not show a dramatic decrease in synaptic density as compared to DA synapses (Figure 2C). However, the integrated synapse intensity was reduced in all neuron types tested, with the most significant decrease in DA neurons (Figure 2D).

In summary, our data suggest that THOC affects overall presynaptic protein levels in all neurons. However, its function is more essential in DA neurons, where it dramatically decreases presynaptic protein level. We propose that there might be a threshold synaptic protein level required for synapse formation. While GABA, serotonin and cholinergic neurons lay above this threshold and form normal number of synapses despite the reduction in presynaptic protein level, DA neurons are below the threshold and hence display a strong reduction in synapse number.

### THOC mutant animals display impaired dopamine signaling and behavioral deficits

To understand whether the morphological synaptic defect leads to a deficit in DA signaling, we analyzed the basal slowing response, a DA dependent behavior (Sawin et al., 2000). Well-fed *wildtype* animals slow down in the presence of food (Figure S2A) and this behavior depends on DA neurons. In contrast, *cat-2* mutants that lack tyrosine hydroxylase and hence cannot synthesize DA, do not slow down on food (Figure S2A). While *thoc-2* heterozygous animals do not show a defect in basal slowing, both *thoc-2* and *thoc-5* homozygous mutants are impaired in this behavior (Figure S2A). Low concentrations of exogenous DA have been shown to restore the basal slowing response in mutants affecting DA synthesis (Figure S2B and Sawin et al., 2000). We tested whether this also holds true for the *thoc-2* and *thoc-5* mutant animals. Indeed, pre-incubating worms on bacterial plates containing 2mM DA restored the basal slowing response back to wild-type (Figure S2B). Taken together, we show that *thoc* mutant animals are highly impaired in DA signaling resulting in a corresponding behavioral deficit.

Impaired basal slowing response can either originate from defective sensory input of the DA neurons, or from defects in signal transmission on the presynaptic site. To rule out that the decreased DA signaling in *thoc* mutant animals arose from sensory defects of the DA neurons, we analyzed the activity of PDE neurons by imaging their calcium transients in freely-behaving worms. Animals expressing the calcium sensor Gcamp6 in DA neurons were imaged during the basal slowing response using an automated calcium imaging system (Hardaway et al., 2015; Piggott et al., 2011). Upon entry into the bacterial food, *wildtype* PDE neurons were activated as measured by an increase in Gcamp6 signal (Figure S2C). Importantly, in *thoc-5* mutant animals, PDEs were equally activated as compared to *wildtype*. These results strongly suggest that defects in DA signaling in *thoc* mutant animals arise from impaired presynaptic development and function. Taken together, we show that *thoc* mutant animals are highly impaired in DA signaling resulting in a corresponding behavioral deficit.

### THOC regulates mRNA export of presynaptic genes

Previous studies implicated the THOC in multiple steps of mRNA maturation, mRNP formation and nuclear export (Luna et al., 2012; Sørensen et al., 2017; Tran et al., 2014a; Wang et al., 2013). In order to elucidate THOC’s role in presynaptic gene expression we performed single molecule in-situ hybridization (smFISH) experiments to specifically visualize and determine the intracellular localization of presynaptic as well as control transcripts in PDE DA neurons. We labeled PDE with GFP for easy cell identification and included the nuclear stain DAPI to delineate the localization of the nucleus. Under *wildtype* conditions, all transcripts predominantly showed cytoplasmic localization with only 10-20% of transcripts retained in the nucleus (Figure 3A, B and Figures S3A). However, in *thoc-5* mutant background, transcripts encoding presynaptic proteins, such as UNC-10/RIM, ELKS-1/ELKS, SYD-2/ liprinα, CLA-1/piccolo and CAT-1/vMAT, displayed 1.5- to 5-fold increase in trapped, nuclear transcripts, while the subcellular localization of control transcripts, including TBA-1/tubulin, UNC-33/CRMP and UNC-44/ankyrin, was unperturbed (Figure 3A, B and Figure S3A).

**Figure 3:**
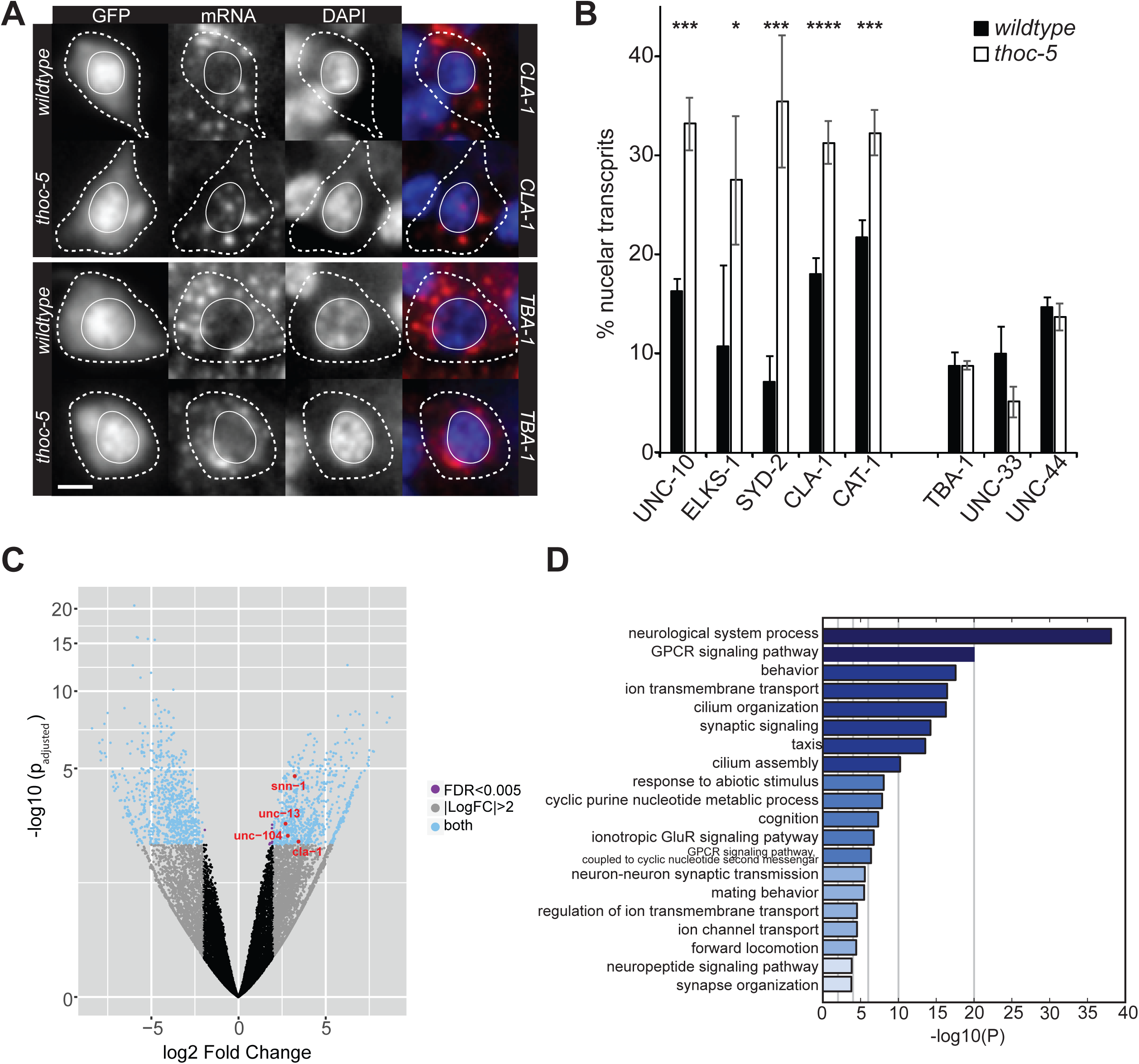
Presynaptic transcripts are trapped in the nucleus in *thoc-5* mutant animals. (A) smFISH for presynaptic (upper panel) and control (lower panel) transcripts in *wildtype* (upper row) and *thoc-5* (lower row) mutant animals. GFP: PDE cellbody; red: signal for FISH probes; blue: DAPI staining of nuclei. Scale bar represents 2μm. (B) Quantification of total transcript levels per PDE neuron in *wildtype* and *thoc-5* mutants. N=10-50, ^****^, p<0.0001; ^***^, p<0.001; ^*^, p<0.05 student t-test. Averages and SEM are plotted in all graphs. (C) Volcano plot of differentially expressed genes between *wildtype* and *thoc-5* mutant neurons. Red genes correspond to significantly up-regulated presynaptic genes further examined throughout this paper. N = 2 biological replicates for *wildtype* and N = 3 biological replicates for *thoc-5* mutant neurons. (D) Gene enrichment analysis for up-regulated genes. See also Figure S3.

Next, we wanted to obtain more comprehensive insight into THOC’s function in neuronal gene expression. To this end, we performed whole transcriptome analysis of *wildtype* and *thoc-5* mutant neurons. Specifically, we dissociated adult worms into single cell suspension through chemo-mechanical disruption, isolated GFP-labeled neurons by fluorescence-activated cell sorting and sequenced their mRNA. We found that 763 genes are up-regulated (fold change ≥ 2 and p < 0.005; 5% of all detected transcripts) and 938 genes are down-regulated (fold change ≤ 2 and p < 0.005, 6% of all detected transcripts) (Figure 3C and Table S1). Gene enrichment analysis revealed that the up-regulated genes are highly enriched in synaptic genes such as synaptic transmission, neurotransmitter receptors and receptor signaling pathways (Figure 3D). Down-regulated genes do not show a strong neuron-specific functional enrichment, they rather associate with a wide variety of biological functions (Figure S3B). Using qPCR, we confirmed the up-regulation of a few presynaptic transcripts in *thoc-5* mutant background (Figure S3C). Interestingly, while we see a decrease in presynaptic proteins, we detect an increase in the corresponding transcript levels. We speculate that this increase in mRNA might be due to a feedback loop, through which the neurons try to boost presynaptic protein production. Alternatively, mRNAs might be trapped in a cellular compartment, where they are inaccessible to translation, however protected from degradation.

Collectively, these observations suggest that THOC plays an important role in the nuclear export and expression of synaptic transcripts, a small, however functionally related set of transcripts specific to neurons.

### THOC5 conditional knock-out mice exhibit loss of dopaminergic synaptic proteins at three weeks of age

To investigate whether the function of THOC5 is evolutionarily conserved in the dopaminergic neurons of mice, we crossed *Thoc5* floxed (*Thoc5fl/fl*) mice with DAT-IRES-Cre mice to produce a conditional knockout of THOC5 in DA neurons (DAT-Cre;*Thoc5*fl/fl, see methods). Interestingly, DAT-Cre;*Thoc5*fl/fl mice exhibited altered growth in body size and locomotion as compared to *wildtype* (Figure 4A, B). THOC5 cKO mice exhibited normal body size and locomotive behavior during the first three postnatal weeks, which was indistinguishable from *wildtype*. However, they started to lose body weight from 4 weeks of age (Figure 4A, B). Furthermore, they became less mobile and acquired hunched body posture by 6 weeks of age. Surprisingly, the compromised body growth and locomotion was accompanied by premature mortality in DAT-Cre;*Thoc5*fl/fl mice at around 6 to 9 weeks of age (Figure 4C). We hypothesize that *thoc5* cKO mice prematurely die due to lack of food intake, as they display severe movement deficits.

**Figure 4.**
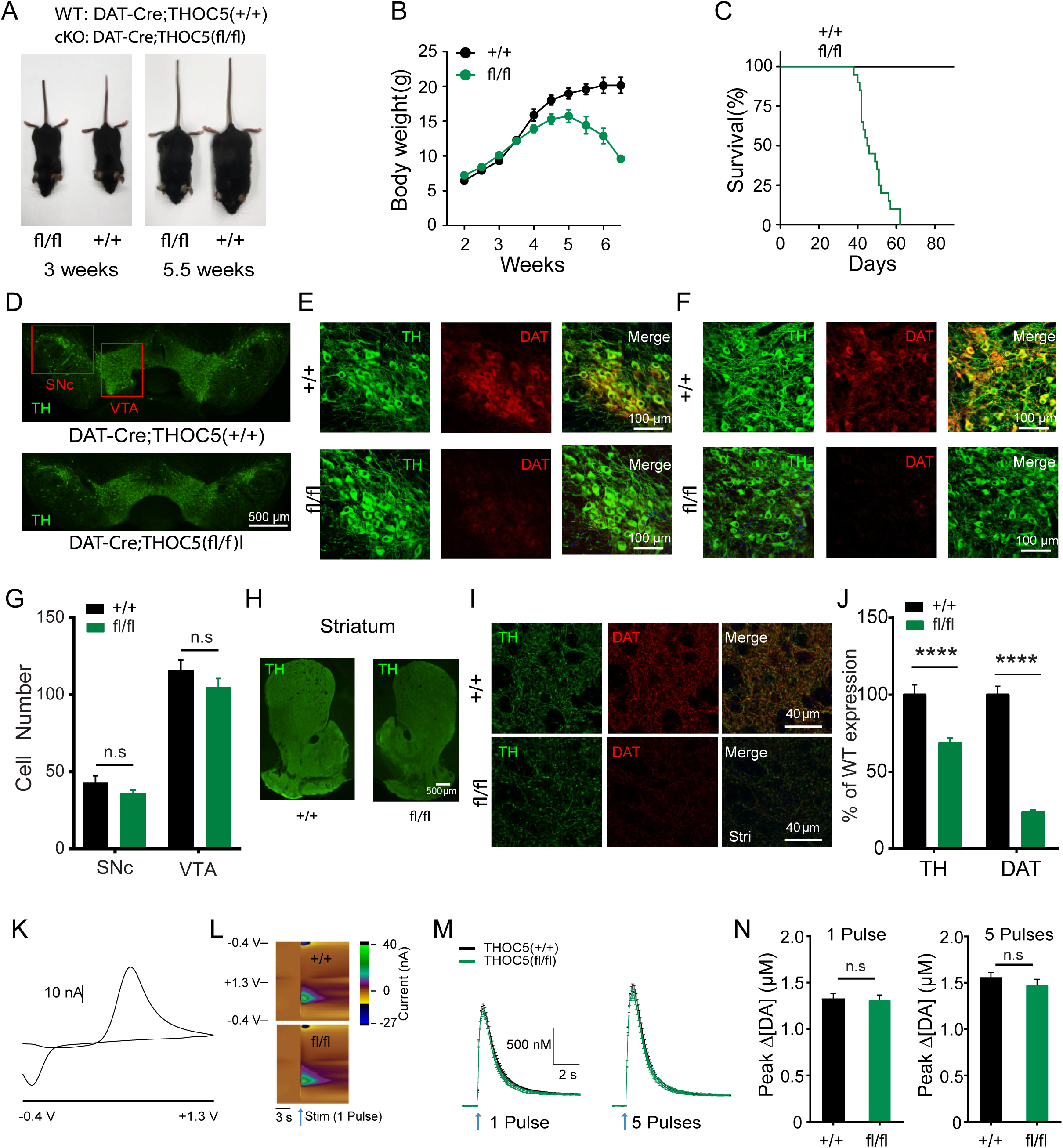
THOC5 conditional knock-out mice exhibit developmental defects, reduced DAT expression, but normal DA release at 3 weeks of age. (A) Representative body size in 3 week old (left) and 5.5 week old (right) DAT-cre;Thoc5+/+ (*wt*) and DAT-cre;Thoc5fl/fl (*cKO*). (B) Summary of *wt* and *cKO* body weight (N=10 for *wt*, and N=10 for *cKO*) (C) Survival curve (N=13 for *wt*, and N=21 for *cKO*) (D) Representative images of midbrain TH immufluorescence in 3 weeks old *wt* and *cKO* mice. (E) Representative images of TH and DAT immunofluorescence in SNc from 3 weeks old *wt* (top) and *cKO* (bottom) mice. (F) Representative images of TH and DAT immunofluorescence in VTA from *wt* (top) and *cKO* (bottom). (G) Quantification of DA cell number in midbrain (p>0.05, two-way ANOVA with Bonferroni post-hoc test, *wt*: N = 12, *cKO*: N = 18) (H) Representative images of striatum showing TH immufluorescence in *wt* (left) and *cKO* (right) mice. (I) Representative high magnification confocal images of TH and DAT immunofluorescence in dorsal striatum from *wt* (top) and *cKO* (bottom). (J) Quantification of TH and DAT immunofluorescence in striatum (^****^, p<0.0001, two-way ANOVA with Bonferroni post-hoc test, *wt*: N = 24, *cKO*: N = 36). (K) 2D voltammogram showing oxidation and reduction peaks of DA. (L) Representative 3D voltammograms from 1 pulse. x-axis: recording time, y-axis: applied potential, color map: recorded current. Arrows represent time of stimulation. (M) Summary of fast-scan cyclic voltammetry (FSCV) recordings measured at the peak oxidative voltage evoked by 1 pulse (left) and 5 pulses (right) stimulations (*wt*: N=24, *cKO*: N=29). (N) Quantification of peak DA release evoked by 1 pulse (left) and 5 pulses (right) stimulations (p>0.05, Mann-Whitney, *wt*: N=24, *cKO*: N=29).

Because DA neurons play an indispensable role in the survival of conditional *thoc5* mutant mice, we next asked whether the conditional deletion of THOC5 has deleterious effects on midbrain DA neurons. To this end, we first checked the number of DA neurons in 3 week-old mice by double immunostaining for tyrosine hydroxylase (TH) and dopamine transporter (DAT) in substantia nigra pars compacta (SNc) and ventral tegmental area (VTA), the cell body region of DA neurons. Confocal imaging of both TH and DAT immunoreactivity demonstrated that there was no difference in DA cell numbers in either SNc or VTA (Figure 4D-G). However, 3 week-old DAT-Cre;*Thoc5*fl/fl mice exhibit less TH, and much less DAT, immunoreactivity in the axon terminals in the striatum, the major target area of midbrain DA neurons (Figure 4H-J).

To assess the physiological consequence of dopaminergic synapse loss in DAT-Cre;*Thoc5*fl/fl mice, we performed fast-scan cyclic voltammetry (FSCV) to measure DA release from dopaminergic axon terminals in the dorsal striatum (Figure 4K-N). We prepared brain slices from these mice and evoked DA release by local electrical stimulation. Consistent with an equal number of TH+ cell bodies in the midbrain, we did not find a significant difference in DA release elicited with 1 or 5 pulses of local electrical stimulation in the dorsal striatum (Figure 4L-N). These data are in agreement with previous studies, where it was shown that depletion of up to 80% of DA neurons was necessary to elicit a deficit in DA release (Zigmond et al., 1984).

Taken together, our data suggest that DA neurons of *thoc5* cKO mice show normal DA neurogenesis and normal axonal outgrowth, but display a substantial loss of presynaptic proteins (DAT) at 3 weeks of age, phenocopying the dramatic synapse loss of DA neurons in THOC mutant worms.

### Dopamine neurons degenerate in THOC5 cKO mice at six weeks of age

To explore whether the synaptic defect of *thoc5* mutant DA neurons aggravated with age as seen for worms (Figure 1F), we conducted the same double immunostaining experiment in ∼6 week-old mice. Confocal imaging of both TH and DAT immunoreactivity demonstrated a significant loss of dopaminergic neurons in DAT-Cre;*Thoc5*fl/fl mice, more severely in SNc than VTA (Figures 5A-D). This phenotype was accompanied by a massive decrease of axon terminals in the striatum (Figure 5E-G). Loss of TH immunoreactivity in dorsal striatum was more pronounced than in ventral striatum, consistent with a greater loss of TH+ cells in SNc compared to VTA.

**Figure 5.**
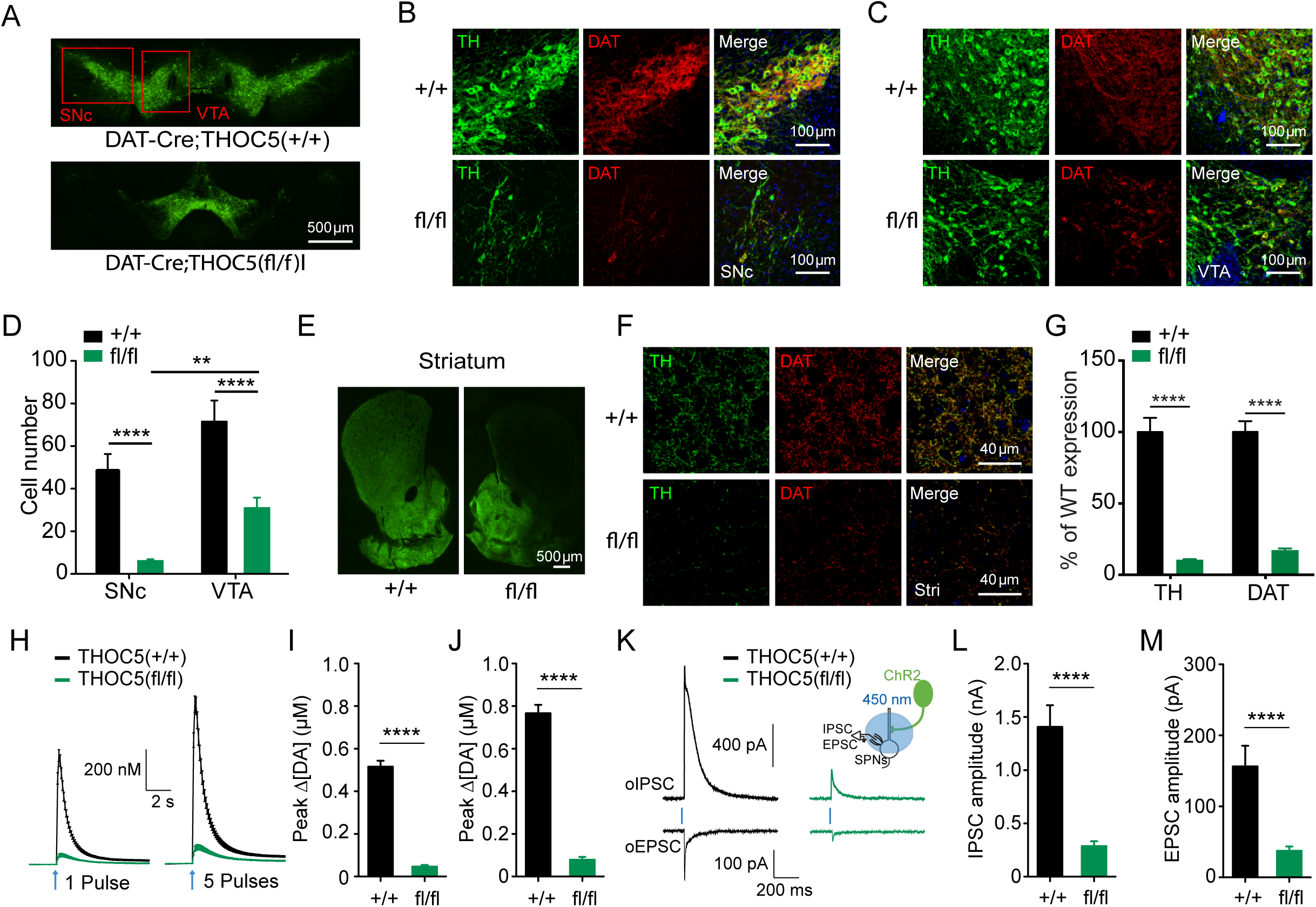
THOC5 conditional knock-out mice exhibit severe loss of DA synapses and diminished DA release at 6 weeks of age. (A) Representative images of midbrain TH immufluorescence in ∼6 week-old DAT-cre;Thoc5+/+ (*wt*) and DAT-cre;Thoc5fl/fl (*cKO*) mice. (B) Representative images of TH and DAT immunofluorescence in SNc from ∼6 week-old *wt* (top) and *cKO* (bottom) mice. (C) Representative images of TH and DAT immunofluorescence in VTA from the same *wt* (top) and *cKO* (bottom) mice. (D) Quantification of DA cell number in midbrain (^**^, p<0.01, ^****^, p<0.0001, two-way ANOVA with Bonferroni post-hoc test, *wt*: N = 10, *cKO*: N = 18) (E) Representative images of striatum showing TH immufluorescence in *wt* (left) and *cKO* (right) mice. (F) Representative high magnification confocal images of TH and DAT immunofluorescence in dorsal striatum from *wt* (top) and *cKO* (bottom) mice. (G) Quantification of TH and DAT immunofluorescence in striatum (^****^, p<0.0001, two-way ANOVA with Bonferroni post-hoc test, *wt*: N = 12, *cKO*: N = 18). (H) Summary of fast-scan cyclic voltammetry (FSCV) recordings measured at the peak oxidative voltage evoked by 1 pulse (left) and 5 pulses (right) stimulations. (I) Quantification of peak DA release evoked by 1 pulse (left) stimulation (^****^, p<0.0001, unpaired t-test, *wt*: N = 13 samples, *cKO*: N = 12). (J) Quantification of peak DA release evoked by 5 pulse (left) 25 Hz stimulation (^****^, p<0.0001, unpaired t-test, *wt*: N = 13 samples, *cKO*: N = 12). (K) Representative traces for oIPSC and oEPSC recorded from *wt* (left) and *cKO* (right) SPNs. Insert: experimental configuration. (L) Quantification of peak oIPSC evoked by optogenetic stimulation (^****^, p<0.0001, unpaired t-test, *wt*: N = 9 samples, *cKO*: N = 14). (M) Quantification of peak oEPSC evoked by optogenetic stimulation (^****^, p<0.0001, unpaired t-test, *wt*: N = 9 samples, *cKO*: N = 13).

To assess the physiological consequence of dopaminergic neuron loss in 6-9 week-old DAT-Cre;*Thoc5*fl/fl mice, we performed FSCV experiments. To selectively activate dopaminergic terminals, DAT-Cre;*Thoc5*fl/fl mice were further bred with Ai32 mice containing a conditional floxed allele of H134R variant of channelrhodopsin2 (ChR2) in the Rosa26 locus, producing DAT-Cre;Ai32;*Thoc5*fl/fl mice. We prepared brain slices from these mice and their *wildtype* controls, and evoked DA release with a short pulse of blue light (1ms, 450 nm, Figure 5H-J). Within the dorsal striatum, DA release from a single light pulse is severely impaired in DAT-Cre;Ai32;*Thoc5*fl/fl mice (Figure 5H-J). To mimic phasic firing of DA neurons, we delivered 5 blue light pulses at 25 Hz; this protocol also revealed a significant deficit (Figure 5H-J). These data suggest that in ∼6 week-old DAT-Cre;*Thoc5*fl/fl mice, DA release is severely diminished, possibly resulting from severe degeneration of DA neuron axons.

There is accumulating evidence that the fast-acting neurotransmitters γ-aminobutyric acid (GABA) and glutamate are co-released along with DA from dopaminergic terminals in the striatum (Hnasko et al., 2010; Kim et al., 2015; Tritsch et al., 2012). These co-released neurotransmitters induce inhibitory postsynaptic currents (IPSC) and much smaller, but reproducible, excitatory postsynaptic currents (EPSC). It is possible that *thoc5* mutant mice have diminished TH and DAT expression but normal synaptic release. To exclude this possibility and validate our finding that dopaminergic axons undergo severe degeneration in ∼6 week-old DAT-Cre;*Thoc5*fl/fl mice, we took advantage of GABA-glutamate co-release from dopaminergic terminals and performed whole-cell patch clamp recordings in postsynaptic striatal spiny projection neurons (SPNs). The amplitude of optogenetically evoked IPSC (oIPSC) and EPSC (oEPSC) are much smaller in DAT-Cre;Ai32;*Thoc5*fl/fl mice as compared to their *wildtype* litter controls (Figure 5K-M).

Together, our immunohistochemical, electrophysiological, and voltammetry data support the conclusion that deletion of *Thoc5* from mouse DA neurons leads to rapid synapse loss followed by DA neuron degeneration and animal death. In conclusion, they unraveled an evolutionary conserved function for THOC in DA synapse function.

### A Human *thoc1* mutation found in ALS patients renders THOC-1 dysfunctional

Defects in nuclear export have become an emerging pathogenic mechanism in many neurodegenerative diseases. As THOC is a nuclear export factor and as our mouse data implicate a potential role for THOC in neurodegeneration, we searched public databases from large human genome-wide association studies for missense and loss-of-function mutations that are found in patients with neurodegenerative diseases, in particular with Parkinson’s disease (PD) and Amyotrophic Lateral Sclerosis (ALS) (Lill et al., 2011, 2012). Multiple missense mutations in THOC1, THOC2 and THOC5 were reported in these databases, however their statistical significance for disease was rather low. To enrich for putative disease causing mutation, we focused on mutations that occurred on conserved amino acid residues and were absent from a much larger dataset of the control human population (Lek et al., 2016). Specifically, we studied the *thoc1* R138H mutation that was found in the ALS database. We introduced this mutation in the corresponding location of the worm *THOC-1* gene, expressed this construct under the DA neuron-specific promoter and assayed, whether it was able to rescue the *thoc* associated dopaminergic synapse impairment. Interestingly, we found that this mutant allele of THOC-1 was unable to rescue the synapse defect (Figure S4). However, the disease-unrelated mutation V338I found in multiple control individuals did not affect the function of THOC-1 and fully rescued the synapse defect in *thoc-1* mutant animals (Figure S4). The R138H mutation lies within the N-terminal domain of the protein, which has been implicated for protein interactions with other THOC components and hence might impact the function of the entire complex (Peña et al., 2012). Taken together, our data shows that the *thoc-1* R138H mutation, which might be associated with ALS, impairs THOC’s function in dopaminergic synaptogenesis.

### THOC’s function in presynaptic gene expression is pivotal under neuronal activity

Recent studies in mouse and *Drosophila* suggest that THOC is not generally needed for bulk mRNA export; rather it facilitates the export of subsets of mRNAs (Guria et al., 2011; Katahira et al., 2009; Rehwinkel et al., 2004; Wang et al., 2013). Furthermore, THOC5 has been shown to regulate the transcription, processing and export of IEGs in various tissues (Tran et al., 2013, 2014b, 2014a). In neurons, IEGs play a prominent role in long-term synaptic changes and memory formation (Okuno, 2011). Many IEGs encode synaptic proteins and their activity-dependent transcription is mediated by the transcription factor CREB.

In order to explore whether neuronal activity modulates THOC-dependent presynaptic protein expression, we silenced PDE activity by a histamine-gated chloride channel (HisCl). Since *C. elegans* does not synthesize histamine, expression of this channel in worm neurons can cause hyperpolarization and silencing of selective neurons upon histamine addition (Pokala et al., 2014). Interestingly, acute silencing of DA neurons by cell-type specific expression of HisCl in *thoc-5* mutant animals partially but significantly restored presynaptic intensity of PDE neurons (Figure 6B). This result suggests that THOC-5 is particularly important for presynaptic protein production when neurons are active. When PDE is silenced, other backup pathways will presumably take over and export the synaptic transcripts out of the nucleus.

**Figure 6:**
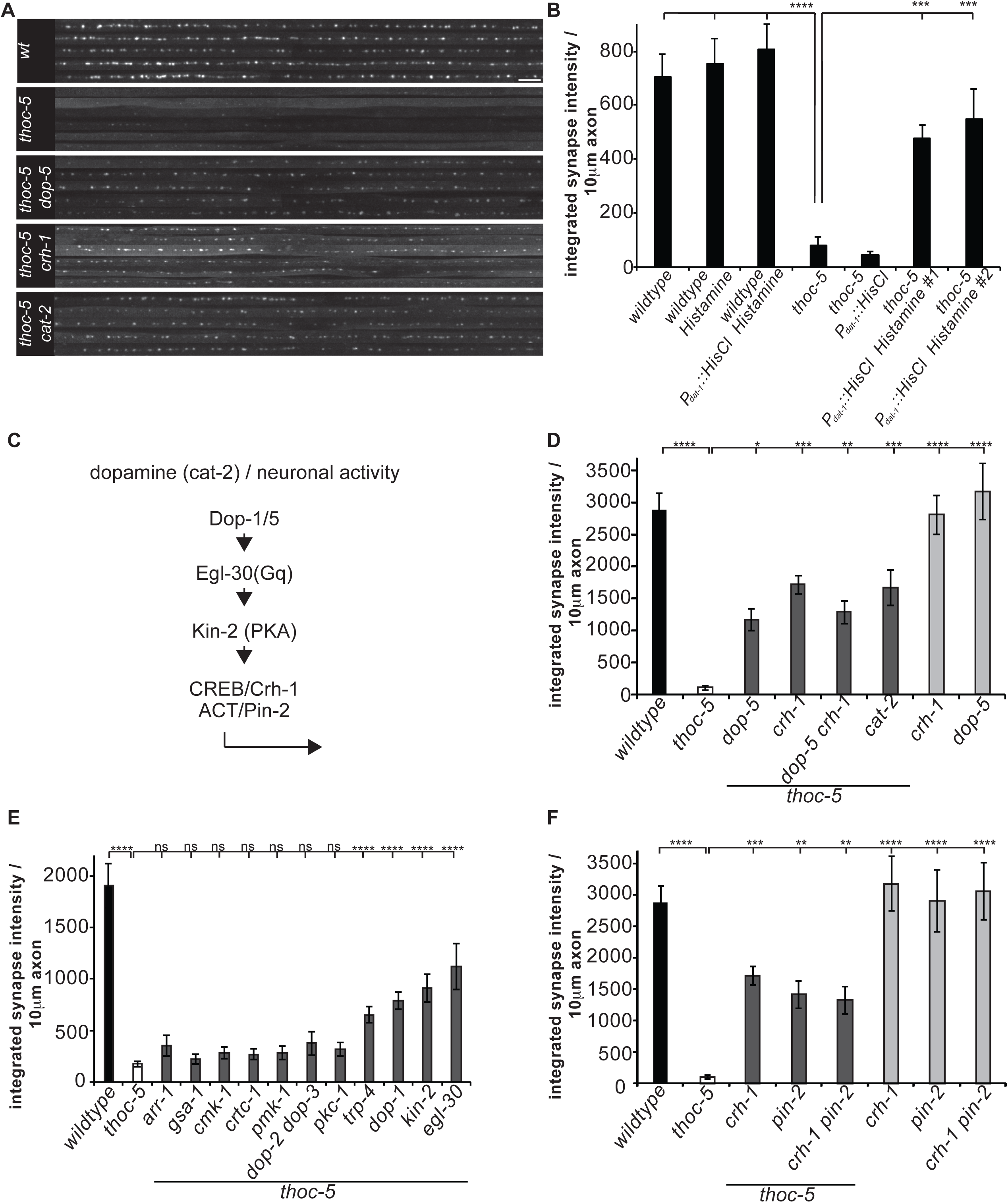
THOC-dependent presynaptic gene expression is pivotal under neuronal activity. (A) Axonal line scans of endogenous GFP::RAB-3 in *wt, thoc-5* and *thoc-5* DA signaling double mutant animals. Scale bar represents 10μm. (B) Silencing of DA neurons by the expression of histamine-gated chloride channel suppresses THOC-dependent synapse defect. Integrated synaptic intensity is quantified. N=10-20 animals; ^****^, p<0.0001; ^***^, p<0.001, one-way ANOVA with Tukey post-hoc test. (C) Schematic of DA signaling cascade feeding into THOC-dependent presynaptic transcription. Neuronal activity modulates THOC-dependent presynaptic transcription through GPCR, trimeric G-protein, PKA and CREB activation. (D) Integrated synaptic intensity is quantified for DA signaling mutants. N=10-20 animals; ^****^, p<0.0001, one-way ANOVA with Tukey post-hoc test. (E) Extensive double mutant analysis identifies signaling components down-stream of neuronal activity. N=10-20 animals; ^****^, p<0.0001, one-way ANOVA with Tukey post-hoc test. (F) CREB functions independently of CBP/CRTC, but in concert with the nuclear factor PIN-2, a LIM-domain containing protein. Integrated synaptic intensity is quantified. N=10-20 animals; ^****^, p<0.0001, one-way ANOVA with Tukey post-hoc test. Averages and SEM are plotted in all graphs. See also Figure S5 & Figure S6.

To further understand the molecular mechanism underlying this activity dependent regulation, we created double mutant strains for *thoc-5* and *crh-1* (*C. elegans* homologue of CREB) and visualized dopaminergic synapses by the endogenously expressed synaptic vesicle marker GFP::RAB-3 (Figure 6A, D). Interestingly, *thoc-5 crh-1* double mutant animals also showed partial but significant rescue of dopaminergic synapses. While synaptic density and size was restored to *wildtype* levels, the intensity of the synapses was only partially rescued (Figure 6A, D). Loss of *crh-1* only in DA neurons of *thoc-5* mutant animals was sufficient to partially restore PDE synapses, suggestive of a cell-autonomous function of CREB in PDE neurons (Figure S5.). *crh-1* single mutants did not show any apparent defect in DA synapses. Interestingly, in other neuronal cell-types, *thoc-5 crh-1* double mutant animals did not show any rescue of presynaptic defects (Figure 2C, D). These observations further underscore THOC’s pivotal role in activity-dependent synaptic gene expression specifically in DA neurons.

As the presynaptic defect was partially rescued *in thoc-5 crh-1* double mutant animals, we wondered whether that was due to a restoration of nuclear export of presynaptic transcripts. Indeed, smFISH experiments in *thoc-5 crh-1* double mutant background revealed that nuclear export of presynaptic transcripts was restored to *wildtype* levels (Figure S6). Together, these data indicate that neuronal activity in PDE induces CREB-mediated transcripts, which are particularly dependent on THOC for nuclear export.

Next, we asked how PDE is activated by examining genetic interactions between *thoc-5* and mutants that affect PDE activity. The *thoc-5* synaptic phenotype could be partially rescued in a *cat-2 thoc-5* double mutant background, which is defective in DA synthesis suggesting that DA itself might activate PDE in an autocrine or paracrine fashion (Figure 6A, D). Consistent with this notion, mutations in DA receptors *dop-1* and *dop-5*, when combined with *thoc-5* loss of function mutation, also mitigated presynaptic loss in PDE indicating that DA actives through these two G-protein coupled receptors (Figure 6A, D & E). In contrast, combining *dop-2* and/or *dop-3* mutations with *thoc-5* did not lead to increased dopaminergic synapses, arguing for receptor specificity (Figure 6E). Furthermore, in a recent publication, the mechanotransduction TRPN/NOMPC channel TRP-4 was shown to be essential for proper DA neuron function and the basal slowing response (Kang et al., 2010). Introducing a mutation in this channel into *thoc-5* mutant strain background partially restored DA synapses (Figure 6E).

To further unravel the signaling cascade involved in linking neuronal activity to CREB in THOC-dependent synapse formation, we performed an extensive double mutant analysis with known upstream signaling molecules of CREB. CREB can be phosphorylated and activated by a large number of kinases, such as PKA, MAP kinases and Ca^2^+/Calmodulin-dependent kinases (CaMKs) (De Cesare et al., 1999). While double mutants between *thoc-5* and the CaMKs *cmk-1* and *unc-43*, the MAPKs *pmk-1* and *pmk-3* and other kinases *pkc-1* and *akt-1* did not show any suppression of dopaminergic synapses (Figure 6E and data not shown), double mutants between *thoc-5* and *kin-2*, the regulatory subunit of PKA, partially restored the synapses (Figure 6E). Furthermore, we identified the heterotrimeric G-protein alpha subunit Gq *egl-30*, but not the Gs *gsa-1* nor beta-arrestin *arr-1* to be involved in this signaling cascade (Figure 6E). CREB often functions in concert with co-activators to promote gene transcription. It is thought that various external stimuli and signaling cascades might lead to recruitment of different co-activators and hence drive expression of specific sets of target genes (Altarejos and Montminy, 2011; Fimia et al., 1999). Double mutant animals between *thoc-5* and *pin-2*, the *C. elegans* homologue of LIM-only protein ACT, but not between *thoc-5* and *crtc-1* showed suppression of synapse loss of the *thoc-5* single mutant animals (Figure 6F).

In conclusion, our data support a strong link between neuronal activity and *thoc-5* dependent presynaptic transcription, which seems to be DA neuron specific. We propose that PDE is activated by DA through *dop-1* and *dop-5* receptors. These receptors in turn activate Gq and PKA to stimulate CREB/ACT dependent transcription. The resulting activity dependent transcripts rely on THOC for efficient nuclear export, subsequent protein translation and synaptogenesis. Hence, in the absence of THOC, the dopaminergic synapse formation program fails.

### CREB interacts with THOC to mark transcripts for efficient export

CREB and THOC were shown to localize to the nucleus, which corresponds well with their function in transcription and mRNA maturation, respectively (Cha-Molstad et al., 2004; Tran et al., 2014b; Wang et al., 2013).

We studied their localization in *C. elegans* DA neurons by expressing fluorescently tagged CREB and THOC proteins under the DA specific promoter. A C-terminal GFP fusion of THOC-2 displayed diffuse localization throughout the cell body, including nucleus and cytoplasm. Furthermore, it accumulated in multiple puncta around the nucleus. We do not know the nature of this compartment, as it did not co-localize with any traditional marker (e.g. Golgi) (Figure 7A and data not shown). THOC-5, on the other hand, was strictly nuclear excluded and showed a diffuse distribution throughout the entire cytoplasm of the neuron (Figure 7A). In non-neuronal cells, THOC-5’s localization varies from mainly nuclear in stem cells and differentiating cells to predominantly cytoplasmic in terminally differentiated macrophages (Tran et al., 2014b). Furthermore, Kamath et al. provide evidence that THOC-5 shuttles between the nucleus and cytoplasm (Kamath et al., 2001). Hence our finding is well in agreement with published data and might reflect THOC-5’s auxiliary role as an adaptor in mRNA export (Katahira et al., 2009).

**Figure 7:**
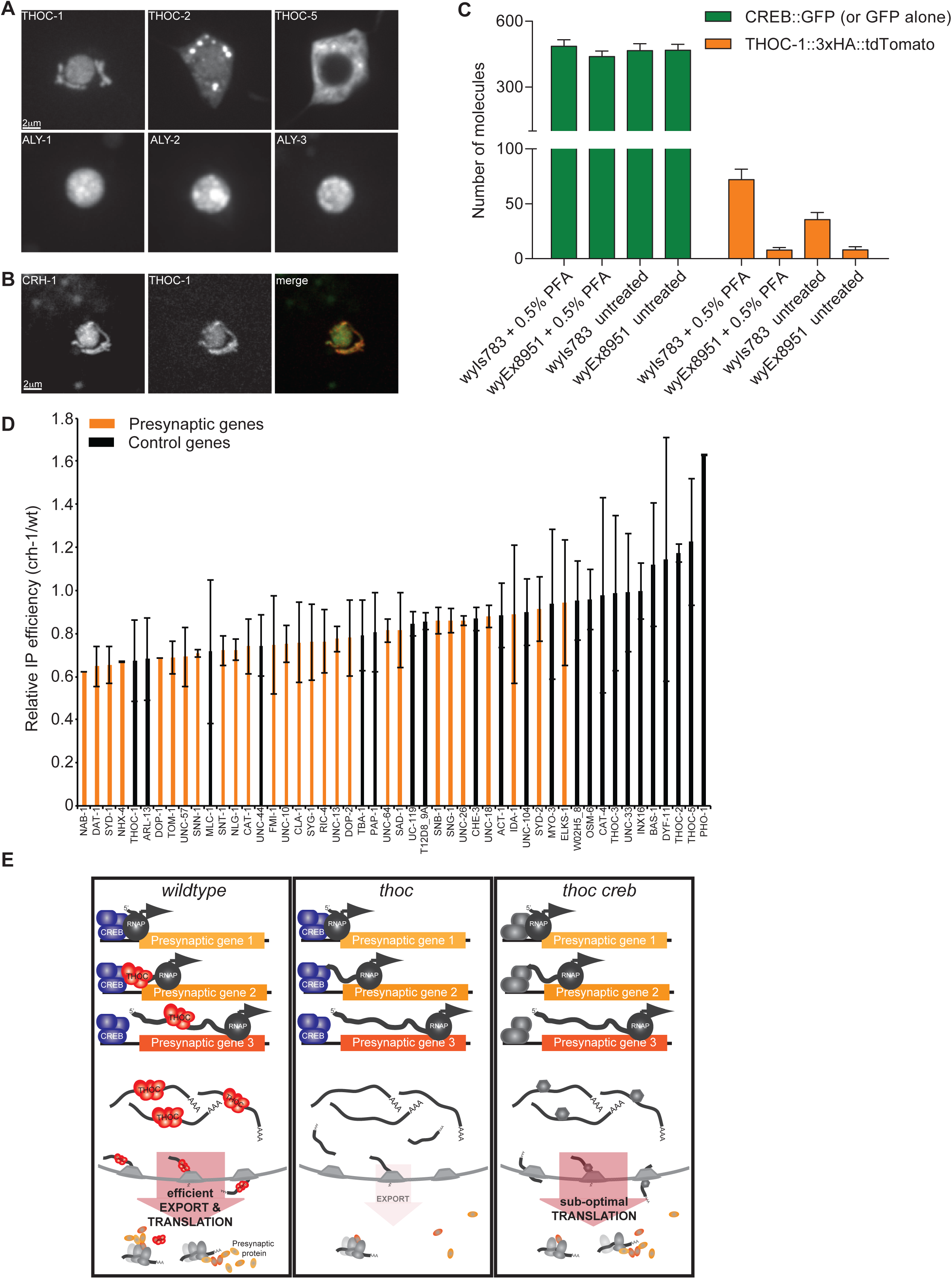
CREB physically interacts with THOC to recruit it onto activity-dependent transcripts. (A) Localization of GFP-tagged THOC components within PDE cell body. Scale bar represents 2□m. (B) Co-expression of fluorescently tagged THOC-1::tdTomato and CREB::GFP show high co-localization in the nucleus and cytoplasmic structures in PDE neurons. Scale bar represents 2□m. (C) Quantification of simPULL assay showing that THOC-1 and CREB transiently interact together. GFP serves as a negative control and does not interact with THOC-1. ^****^, p<0.0001 student t-test. (D) Immunoprecipitation of THOC-1-bound RNAs from *wildtype* and *crh-1* mutant neurons. Immunoprecipitated RNAs were quantified by qPCR. Many presynaptic genes (orange) are less efficiently precipitated in *crh-1* mutant background as compared to *wildtype* conditions. Other pan-neuronal and control genes (black) do not depend as much on CREB. Averages and SEM are plotted in all graphs. (E) Current model of how THOC functions to regulate presynaptic gene expression in DA neurons. Upper panel represents *wildtype*, middle panel *thoc* single and lower panel *thoc creb* double mutant situation. In *wildtype*, CREB recruits THOC onto presynaptic transcripts to facilitate their efficient export into the cytoplam for their concerted translation. In *thoc* mutant background, these transcripts get trapped in the nucleus, hence leading to decreased presynaptic proteins and defective synapse assembly. This defect can be alleviated by removal of CREB, thereby allowing for a constitutive backup export pathway to kick in and to re-establish efficient export of presynaptic transcripts. See also Figure S7.

The three AlyRef/THOC-4 homologues ALY-1,-2,-3, THOC-1 and CREB showed predominant nuclear localization (Figure 7A, B). Interestingly, THOC-1 and CREB also localized to an additional compartment in the cytoplasm. We speculate that this compartment represents mitochondria, as CREB was shown to localize there (Sepuri et al., 2016). Co-expression of THOC-1 and CREB in PDE neurons revealed perfect co-localization of the two proteins (Figure 7B). Collectively, our localization data supports our genetic link between THOC and CREB and raises the question whether they might work together through physical interaction to facilitate presynaptic gene expression.

To address whether THOC and CREB directly interact with each other, we performed single molecule pull-down (SiMPull) experiments. SiMPull is an imaging-based, quantitative co-immunoprecipitation assay that allows to detect weak and transient binding due to its high sensitivity (Jain et al., 2011). We constructed strains expressing THOC-1::3xHA::tdTomato with either CREB::GFP (wyIs783) or GFP (wyEx8951) under a pan-neuronal promoter. As we assumed that the interaction between THOC and CREB might be transient, we also crosslinked one batch of worms with paraformaldehyde. Worm lysates were then subjected to co-immunoprecipitation with a biotinylated anti-GFP antibody, and the pull-down fractions were analyzed by fluorescence microscopy. While THOC-1 did not bind to GFP alone in neither the untreated nor the fixed sample (Figure 7C, Figure S7A), we detected a small, but significant number of THOC-1 molecules interacting with CREB under native condition. Furthermore, the detected interaction was more prominent when samples were fixed with paraformaldehyde (Figure 7C). Taken together our data supports a model, where transient interaction between CREB and THOC-1 might mark presynaptic transcript for efficient export out of the nucleus.

To directly test this model, we performed RNA immunoprecipitation experiments. Specifically, we immunoprecipitated pan-neuronally expressed THOC-1::3xHA::tdTomato under *wildtype* and CREB mutant conditions and quantified the THOC-1-bound and co-immunoprecipitated mRNAs by qPCR. We hypothesized that in a CREB mutant strain background, THOC would be less efficiently loaded onto activity-dependent transcripts and hence, less activity-dependent mRNAs would be precipitated. In line with our hypothesis, we found that THOC-1 bound considerable amounts of presynaptic transcripts under *wildtype* conditions (Figure S7B). Under CREB mutant conditions, roughly 2-fold less mRNA was precipitated by THOC-1 as compared to *wildtype*. Furthermore, precipitation of presynaptic transcripts was strongly dependent on the presence of CREB, while other neuronal transcripts and housekeeping transcripts were less affected (Figure 7D). In conclusion we propose a model, where activity-dependent transcripts get efficiently exported out of the nucleus due to the CREB-dependent loading of THOC onto these transcripts (Figure 7E).

## Discussion

During neuronal differentiation, the majority of synapses in a given neuron form in a relatively short period of time, during which the expression of synaptic proteins needs to be up-regulated in a concerted manner. How the synthesis of synaptic proteins is coordinated, is not well understood. Through an unbiased forward genetic screen, we identified the THOC as a master regulator of presynaptic gene expression especially in DA neurons. Loss of THOC leads to dramatic synapse development deficit in both worms and mouse. THOC is recruited onto activity-dependent mRNAs by direct physical interaction with the CREB transcription factor, thereby facilitating efficient nuclear export and translation of presynaptic transcripts (Figure 7E). Taken together, these results reveal another layer of neuronal gene expression regulation and demonstrate that nuclear export is an important regulatory mechanism for neurodevelopment.

### Transcription factor-dependent loading of THOC onto mRNAs as a novel mechanism for target selectivity

THOC was shown to be essential during early mouse development, as *thoc5* and *thoc1* full knock-out animals die during embryogenesis (Mancini et al., 2010; Wang et al., 2006). Furthermore, this protein complex is also indispensable for survival of adult mice, as full knock-out of *thoc5* after birth leads to rapid death. Interestingly, even though THOC seems to be a very important RNA metabolism factor, it does not affect mRNA on a global level, rather it was shown to regulate subsets of transcripts in different tissues and under diverse conditions. In *Drosophila*, less than 12% of total transcripts are affected by THOC depletion (Rehwinkel et al., 2004). In Hela cells, mouse embryonic fibroblasts, embryonic stem cells and macrophages THOC knockdown led to the down-regulation of 289, 228, 143 and 99 genes respectively (Guria et al., 2011; Katahira et al., 2013; Tran et al., 2013; Wang et al., 2013). How THOC selects its targets is still an open question in the field. In yeast, two studies have shown that THOC is required for the transcription of long and/or GC rich transcripts as well as tandem repeat sequences (Chávez et al., 2001; Voynov et al., 2006). While some mammalian THOC targets contain higher GC content or repeat sequences, these two features are not sufficient to specify THOC targets. In higher model organisms, most known THOC-dependent transcripts contain introns (Guria et al., 2011; Saran et al., 2013; Tran et al., 2013), however a direct link for THOC in splicing has only been established in plants (Sørensen et al., 2017). In mammalian cell culture, THOC5 was shown to paly a role in 3’ end processing through direct interaction with different members of the cleavage/polyadenylation complex (Katahira et al., 2013; Tran et al., 2014a). As most genes in metazoans contain introns and multiple polyadenylation signals, these mechanisms can’t explain THOC’s target selectivity satisfyingly. Here we provide evidence for a novel target selectivity mechanism for THOC. Our data supports a model, whereby direct interaction between a transcription factor and the THOC recruits this complex directly onto nascent transcripts, thereby selectively marking them for efficient downstream processing and export out of the nucleus. This model offers an explanation on how the THOC RNA binding complex with thus far unknown and “unspecific” RNA binding affinities (e.g. Wang et al. showed that THOC2 can bind throughout the entire spliced mRNA of its targets) can still selectively bind to a specific group of target mRNAs. Furthermore, this model provides an organism with an elegant mechanism to harness a single RNA binding complex to facilitate distinct gene expression profiles by coupling different transcription factors to THOC and thereby providing an express lane for certain mRNAs out of the nucleus for efficient downstream protein translation.

### THOC and CREB work together to efficiently export activity-induced transcripts

In our study, we describe a novel function of THOC in terminally differentiated neurons. Our understanding of THOCs function in the brain so far only based on human genetic studies (Amos et al., 2017; Anazi et al., 2016; Beaulieu et al., 2013; Boycott et al., 2010; Di Gregorio et al., 2013; Kumar et al., 2015), however how THOC affects neurons at the cellular or molecular level is completely unknown. We show that THOC does not affect neuronal proliferation and neuron fate determination, but rather has a conserved role in synapse formation and maintenance. In THOC mutant worms and mice, DA neurons proliferate and develop normally, however dopaminergic synapses are dramatically reduced. Our mRNA localization studies in THOC mutant worms revealed a nuclear export defect specifically for presynaptic, but not control transcripts. How does THOC affect a specific set of genes including presynaptic genes? First clues came from studies in non-neuronal cells, where it was shown that THOC has a prominent role in the processing and export of IEGs in LPS-stimulated macrophages and serum-stimulated MEFs (Tran et al., 2013, 2014a). In neurons, IEGs are induced by neuronal activity and play an important role in synaptic plasticity and memory formation. Interestingly, many synaptic genes can be classified as IEGs in neurons (Impey et al., 2004; Lakhina et al., 2015; Okuno, 2011). In this study we propose a model, whereby the activity-dependent transcription factor CREB loads THOC specifically onto activity-induced transcripts (Figure 7E). Through genetics and biochemistry, we show that CREB directly interacts with THOC (Figure 6A, D and Figure 7C) and that CREB is necessary to efficiently recruit THOC onto synaptic transcripts (Figure 7D). The activity dependent, CREB-mediated transcripts are particularly dependent on THOC, which is demonstrated by the striking loss of multiple presynaptic proteins in the THOC single mutants. Interestingly, under the conditions of reduced neuronal activity or failed CREB activation, the compromised nuclear export of mRNAs and concomitant reduced protein translation is partially rescued, suggesting that other export pathways, probably more constitutive backup pathways, can partially substitute for THOC-dependent export. However, these backup pathways are not capable of fully replacing the THOC-dependent mechanisms, as DA synapses are significantly dimmer in *thoc-5 crh-1* double mutant animals (Figure 6A, D). Since the smFISH experiments in *thoc-5 crh-1* double mutant background revealed that the nuclear export of presynaptic transcripts was restored to *wildtype* levels in DA neurons (Figure S3B), we suspect that THOC affects other aspects of the mRNA that impact their optimal translational activity. For example, Tran et al showed, that some IEG transcripts were differently 3’end cleaved in the absence of *thoc5* (Tran et al., 2014a).

### A conserved role for THOC in dopaminergic synaptogenesis

As THOC and its molecular function in mRNA maturation and export is conserved from yeast to mammals, and as multiple human genetic studies implicated THOC in normal brain function, we explored whether THOC’s role in presynaptic gene expression, as discovered in *C. elegans*, was conserved in mice. Indeed, conditional knock-out of *thoc5* in mouse DA neurons led to a significant loss of presynaptic proteins at 3 weeks of age, while DA neuron number was unchanged (Figure 4). These data nicely recapitulate the worm *thoc-5* mutant phenotype. Furthermore, they underscore THOC’s new function in terminally differentiated cells. *thoc5* conditional knock-out mice at 6 weeks of age displayed an even more dramatic synapse loss due to DA neuron degeneration, which resulted in ataxia and eventual animal death (Figure 4 and 5). While we could not detect DA neuron degeneration and cell death in *C. elegans*, we indeed found an age-dependent decline of DA synapses, mirroring the progressive synapse loss in mice (Figure 1F). We speculate that *C. elegans* do not live long enough to exhibit dramatic dopaminergic degeneration as seen in mice.

Loss of DA neurons in *thoc5* conditional mutant mice could be attributed to an essential, house-keeping function of THOC in these neurons. However, we believe that this is rather unlikely. First, essential genes often impact cell growth and proliferation. *thoc-5* mutant mice however contained the same number of DA neurons as their *wildtype* littermates at three weeks of age, when the mouse DA neuron system is fully mature (Prakash and Wurst, 2006). Second, loss of *thoc5* differentially affected DA neuron degeneration in the midbrain. SNc DA neurons displayed a much stronger degenerative loss as compared to their VTA counterparts (Figure 4). Third, multiple mouse studies (Tran et al., 2014b; Wang et al., 2013) as well as our *C. elegans* study showed that THOC did not affect overall protein expression, but rather affected the expression of a small subsets of genes.

Might loss of THOC paly a role in neurodegenerative disease? While defects in nuclear export has emerged as a common cause of multiple neurodegenerative diseases (Freibaum et al., 2015; Tsoi et al., 2011; Zhang et al., 2015), and components of the THOC were shown to mislocalize in these diseases (Woerner et al., 2016), the causative link between THOC mislocalization and neurodegeneration still awaits further investigation. Interestingly, in our mouse work, we found that DA neurons of the SNc degenerated much more dramatically as compared to the ones of the VTA, a phenomenon reminiscent of PD. Furthermore, it is noteworthy that large human genome-wide association studies identified multiple mis-sense and loss-of-function mutations in different THOC components that show association with neurodegenerative diseases, in particular with ALS and to a lesser extent with PD (Lill et al., 2011, 2012). Introduction of the disease associated *thoc1* R138H mutation in the corresponding amino acid of the worm THOC-1 protein rendered the mutant protein dysfunctional, such that it was unable the rescue the synapse defect of worm DA neurons (Figure S4). A disease-unrelated mutation found in multiple control individuals (Lek et al., 2016) however did not affect the function of THOC-1 and fully rescued the synapse defect in *thoc-1* mutant animals. These data might offer a first glimpse into THOC potential role in neurodegeneration. In the future, further massive sequencing efforts of human genomes, together with continuing studies in mouse and *C. elegans* will be needed to shed more light onto THOC’s potential role in neurodegeneration.

## Author Contributions

C.I.M. and K.S. conceived, conducted and analyzed the experiments and wrote the paper. J.I.K., K.K. and J.B.D. did all the mouse experiments, J.B.D. contributed to the manuscript writing. A.S. and Y.K.X. did the simPull experiment. Z.L. and X.Z.S.X performed calcium imaging. Q.L and J.B.L. performed the RNAseq analysis.

## Acknowledgements

We are grateful to Michael L. Nonet for sharing antibodies against *C.elegans* presynaptic proteins. We thank the international Caenorhabditis Genetics Center for strains. We are grateful to Teruko Tamura for providing us with the floxed THOC5 mice. We also thank C. Gao and K. Vega for technical assistance and members of the Shen lab for valuable comments and discussion. This work is supported by the Howard Hughes Medical Institute and the NIH (K.S.)

---

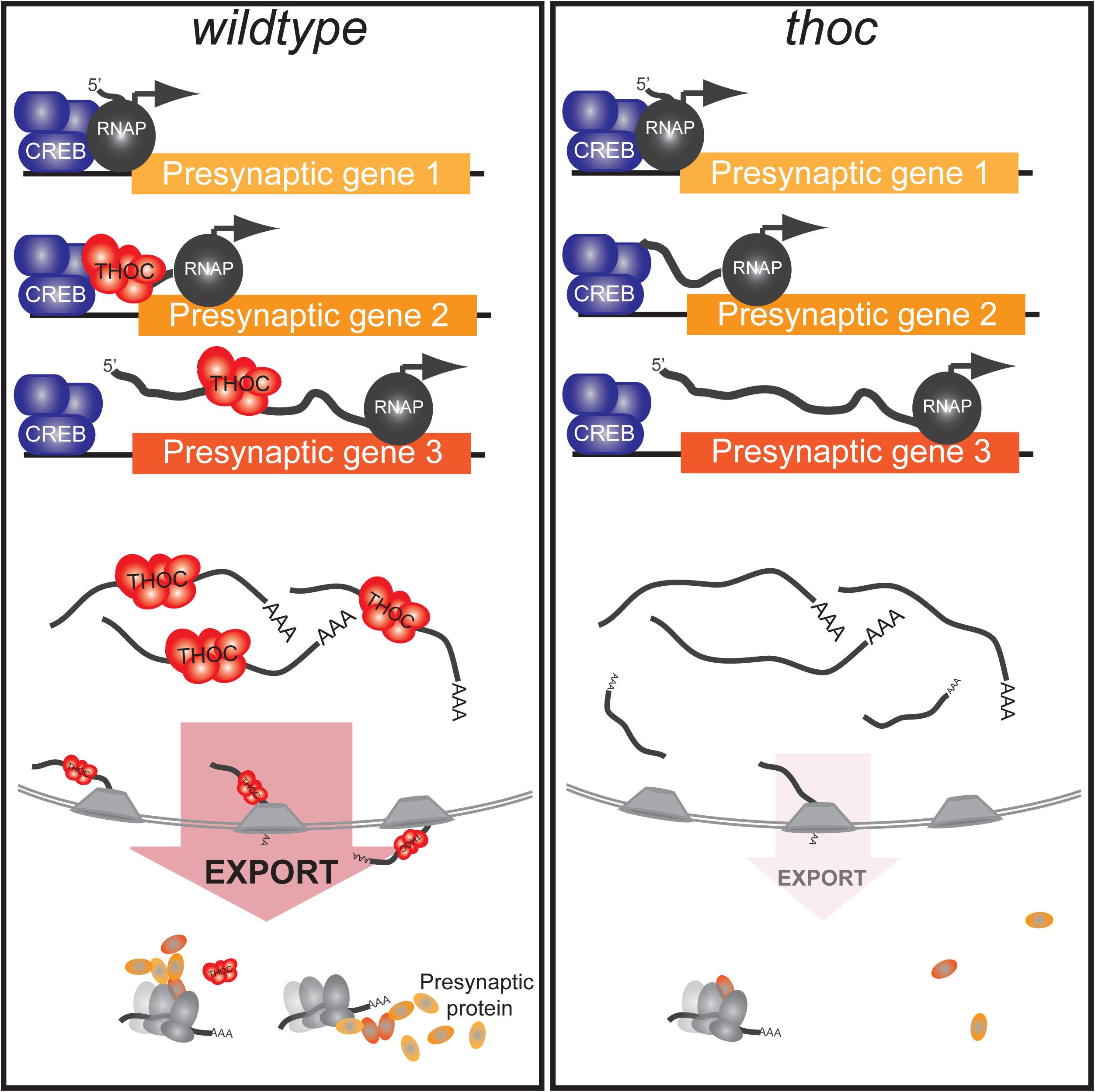

**Figure S1: Detailed analysis of dopaminergic synapse defects in THOC mutant animals**

(A) Quantification of synaptic density relative to wildtype for different THOC mutants and their corresponding cell-specific rescue.

(B) Quantification of synaptic size for different THOC mutants and their corresponding cell-specific rescue.

(C) Quantification of synapse intensity for different THOC mutants and their corresponding cell-specific rescue.

(A-C) N=10-20 animals at L4 stage; ^****^, p<0.0001, ^***^, p<0.001^**^, p<0.01, ^*^, p<0.05, one-way ANOVA with Tukey post-hoc test.

(D) Quantification of synaptic density for L3 and L4 animals in *wildtype* and *thoc-5* mutant background.

(E) Quantification of synaptic size for L3 and L4 animals in *wildtype* and *thoc-5* mutant background.

(F) Quantification of synapse intensity for L3 and L4 animals in *wildtype* and *thoc-5* mutant background.

(D-F) N=10-20 animals; ^***^, p<0.001, student t-test.

Averages and SEM are plotted in all graphes.

**Figure S2: THOC mutants are impaired in DA signaling and DA-dependent behavior**

(A) and (B) Quantitative analysis of basal slowing response in the absence (A) or presence (B) of DA. Locomotion rates in the absence (black bars) or presence (white bars) of bacteria were calculated in a 20 second time window. *Wildtype* and *cat-2* animals were included as controls. The percentage of slowing is shown on the right. N=30-60 animals; ^***^, p<0.001, ^**^, p<0.01, one-way ANOVA with Tukey post-hoc test. (C) Ca^2^+ response of PDE neuron during basal slowing response. Calcium trace and quantification of it is shown for *wildtype* and *thoc-5* mutant animals.

Averages and SEM are plotted in all graphs.

**Figure S3: THOC specifically regulates synaptic transcripts as analyzed by smFISH, RNAseq and qPCR**

(A) Representative images for smFISH for presynaptic and control transcripts in *wildtype* (upper row) and *thoc-5* (lower row) mutant animals. GFP: PDE cellbody; red: signal for FISH probes; blue: DAPI staining of nuclei. Scale bar represents 2μm.

(B) Gene enrichment analysis for down-regulated genes in *thoc-5* mutant neurons.

(C) qPCR quantification of presynaptic as well control transcripts in *wildtype* and *thoc-5* mutant strain backgrounds. N = 3 biological replicates for both genotypes. ^****^, p<0.0001, ^***^, p<0.001, ^**^, p<0.01; one-way ANOVA with Tukey post-hoc test. Averages and standard deviations are plotted.

**Figure S4: Putative disease-associated *thoc-1* R123H mutation renders THOC-1 dysfunctional**

Integrated synaptic intensity is quantified for *wt, thoc-1, thoc-1 P*_*dat-1*_*::thoc-1R123H* and *thoc-1 P*_*dat-1*_*::thoc-1V338I* strain backgrounds. N=20 animals; ^****^, p<0.0001, one-way ANOVA with Tukey post-hoc test.

Averages and SEM are plotted.

**Figure S5: CREB functions cell-autonomously in DA neurons to suppress *thoc*-dependent presynaptic defects**

(A) Integrated synaptic intensity is quantified for two lines, in which *crh-1* is cell-specifically knocked-down in DA neuron by CRE-loxP strategy. N=10-20 animals; ^****^, p<0.0001, ^***^, p<0.001, ^**^, p<0.01 one-way ANOVA with Tukey post-hoc test.

**Figure S6: Presynaptic transcripts are exported out of the nucleus in *thoc-5 creb* mutant animals**

Quantification of nuclear transcripts in *wildtype, thoc-5, thoc-5 crh-1* and *crh-1* PDE neurons. N=10-50 animals. ^***^, p<0.001; one-way ANOVA with Tukey post-hoc test. Averages and SEM are plotted.

**Figure S7: THOC-1 specifically interacts with CREB but not GFP and binds to presynaptic transcripts**

(A) Quantification of the interaction between THOC-1 and increasing GFP concentration. No significant interaction between those two proteins was detected even at very high GFP concentration in PFA fixed samples. These result further corroborate the specificity of the direct interaction between CREB and THOC-1.

(B) Relative IP efficiency of presynaptic transcripts (orange bars) as well as other pan-neuronal and control genes (black). THOC-1 bound transcripts were pulled-down under *wildtype* conditions. Transcripts were then quantified by qPCR and normalized onto total mRNA.

## References

Altarejos, J.Y., and Montminy, M. (2011). CREB and the CRTC co-activators: sensors for hormonal and metabolic signals. Nat. Rev. Mol. Cell Biol. 12, 141–151.

Amos, J. s., Huang, L., Thevenon, J., Kariminedjad, A., Beaulieu, C. l., Masurel-Paulet, A., Najmabadi, H., Fattahi, Z., Beheshtian, M., Tonekaboni, S. h., et al. (2017). Autosomal recessive mutations in THOC6 cause intellectual disability: syndrome delineation requiring forward and reverse phenotyping. Clin. Genet. 91, 92–99.

Anazi, S., Alshammari, M., Moneis, D., Abouelhoda, M., Ibrahim, N., and Alkuraya, F.S. (2016). Confirming the candidacy of THOC6 in the etiology of intellectual disability. Am. J. Med. Genet. A. 170, 1367–1369.

Arroyo, D.A., and Feller, M.B. (2016). Spatiotemporal Features of Retinal Waves Instruct the Wiring of the Visual Circuitry. Front. Neural Circuits 10.

Bartsch, D., Casadio, A., Karl, K.A., Serodio, P., and Kandel, E.R. (1998). CREB1 Encodes a Nuclear Activator, a Repressor, and a Cytoplasmic Modulator that Form a Regulatory Unit Critical for Long-Term Facilitation. Cell 95, 211–223.

Beaulieu, C.L., Huang, L., Innes, A.M., Akimenko, M.-A., Puffenberger, E.G., Schwartz, C., Jerry, P., Ober, C., Hegele, R.A., McLeod, D.R., et al. (2013). Intellectual disability associated with a homozygous missense mutation in THOC6. Orphanet J. Rare Dis. 8, 62.

Botti, V., McNicoll, F., Steiner, M.C., Richter, F.M., Solovyeva, A., Wegener, M., Schwich, O.D., Poser, I., Zarnack, K., Wittig, I., et al. (2017). Cellular differentiation state modulates the mRNA export activity of SR proteins. J Cell Biol jcb.201610051.

Boycott, K.M., Beaulieu, C., Puffenberger, E.G., McLeod, D.R., Parboosingh, J.S., and Innes, A.M. (2010). A novel autosomal recessive malformation syndrome associated with developmental delay and distinctive facies maps to 16ptel in the Hutterite population. Am. J. Med. Genet. A. 152A, 1349–1356.

Cha-Molstad, H., Keller, D.M., Yochum, G.S., Impey, S., and Goodman, R.H. (2004). Cell-type-specific binding of the transcription factor CREB to the cAMP-response element. Proc. Natl. Acad. Sci. U. S. A. 101, 13572–13577.

Chávez, S., Beilharz, T., Rondón, A.G., Erdjument-Bromage, H., Tempst, P., Svejstrup, J.Q., Lithgow, T., and Aguilera, A. (2000). A protein complex containing Tho2, Hpr1, Mft1 and a novel protein, Thp2, connects transcription elongation with mitotic recombination in Saccharomyces cerevisiae. EMBO J. 19, 5824–5834.

Chávez, S., García-Rubio, M., Prado, F., and Aguilera, A. (2001). Hpr1 Is Preferentially Required for Transcription of Either Long or G+C-Rich DNA Sequences in Saccharomyces cerevisiae. Mol. Cell. Biol. 21, 7054–7064.

De Cesare, D., Fimia, G.M., and Sassone-Corsi, P. (1999). Signaling routes to CREM and CREB: plasticity in transcriptional activation. Trends Biochem. Sci. 24, 281–285.

Di Gregorio, E., Bianchi, F.T., Schiavi, A., Chiotto, A.M.A., Rolando, M., Verdun di Cantogno, L., Grosso, E., Cavalieri, S., Calcia, A., Lacerenza, D., et al. (2013). A de novo X;8 translocation creates a PTK2-THOC2 gene fusion with THOC2 expression knockdown in a patient with psychomotor retardation and congenital cerebellar hypoplasia. J. Med. Genet. 50, 543–551.

Fimia, G.M., De Cesare, D., and Sassone-Corsi, P. (1999). CBP-independent activation of CREM and CREB by the LIM-only protein ACT. Nature 398, 165–169.

Frank, R.A., and Grant, S.G. (2017). Supramolecular organization of NMDA receptors and the postsynaptic density. Curr. Opin. Neurobiol. 45, 139–147.

Freibaum, B.D., Lu, Y., Lopez-Gonzalez, R., Kim, N.C., Almeida, S., Lee, K.-H., Badders, N., Valentine, M., Miller, B.L., Wong, P.C., et al. (2015). GGGGCC repeat expansion in C9orf72 compromises nucleocytoplasmic transport. Nature 525, 129–133.

Guria, A., Tran, D.D.H., Ramachandran, S., Koch, A., Bounkari, O.E., Dutta, P., Hauser, H., and Tamura, T. (2011). Identification of mRNAs that are spliced but not exported to the cytoplasm in the absence of THOC5 in mouse embryo fibroblasts. RNA 17, 1048–1056.

Hardaway, J.A., Sturgeon, S.M., Snarrenberg, C.L., Li, Z., Xu, X.Z.S., Bermingham, D.P., Odiase, P., Spencer, W.C., Miller, D.M., Carvelli, L., et al. (2015). Glial Expression of the Caenorhabditis elegans Gene swip-10 Supports Glutamate Dependent Control of Extrasynaptic Dopamine Signaling. J. Neurosci. 35, 9409–9423.

Heath, C.G., Viphakone, N., and Wilson, S.A. (2016). The role of TREX in gene expression and disease. Biochem. J. 473, 2911–2935.

Hnasko, T.S., Chuhma, N., Zhang, H., Goh, G.Y., Sulzer, D., Palmiter, R.D., Rayport, S., and Edwards, R.H. (2010). Vesicular Glutamate Transport Promotes Dopamine Storage and Glutamate Corelease In Vivo. Neuron 65, 643–656.

Hobert, O. (2016). Terminal Selectors of Neuronal Identity. Curr. Top. Dev. Biol. 116, 455–475.

Iacoangeli, A., and Tiedge, H. (2013). Translational control at the synapse: role of RNA regulators. Trends Biochem. Sci. 38, 47–55.

Impey, S., McCorkle, S.R., Cha-Molstad, H., Dwyer, J.M., Yochum, G.S., Boss, J.M., McWeeney, S., Dunn, J.J., Mandel, G., and Goodman, R.H. (2004). Defining the CREB Regulon: A Genome-Wide Analysis of Transcription Factor Regulatory Regions. Cell 119, 1041–1054.

Jain, A., Liu, R., Ramani, B., Arauz, E., Ishitsuka, Y., Ragunathan, K., Park, J., Chen, J., Xiang, Y.K., and Ha, T. (2011). Probing cellular protein complexes using single-molecule pull-down. Nature 473, 484–488.

Jimeno, S., Rondón, A.G., Luna, R., and Aguilera, A. (2002). The yeast THO complex and mRNA export factors link RNA metabolism with transcription and genome instability. EMBO J. 21, 3526–3535.

Kamath, R.V., Leary, D.J., and Huang, S. (2001). Nucleocytoplasmic Shuttling of Polypyrimidine Tract-binding Protein Is Uncoupled from RNA Export. Mol. Biol. Cell 12, 3808–3820.

Kang, L., Gao, J., Schafer, W.R., Xie, Z., and Xu, X.Z.S. (2010). C. elegans TRP Family Protein TRP-4 Is a Pore-Forming Subunit of a Native Mechanotransduction Channel. Neuron 67, 381–391.

Katahira, J., Inoue, H., Hurt, E., and Yoneda, Y. (2009). Adaptor Aly and co-adaptor Thoc5 function in the Tap-p15-mediated nuclear export of HSP70 mRNA. EMBO J. 28, 556–567.

Katahira, J., Okuzaki, D., Inoue, H., Yoneda, Y., Maehara, K., and Ohkawa, Y. (2013). Human TREX component Thoc5 affects alternative polyadenylation site choice by recruiting mammalian cleavage factor I. Nucleic Acids Res. 41, 7060–7072.

Kauffman, A.L., Ashraf, J.M., Corces-Zimmerman, M.R., Landis, J.N., and Murphy, C.T. (2010). Insulin Signaling and Dietary Restriction Differentially Influence the Decline of Learning and Memory with Age. PLoS Biol. 8, e1000372.

Kim, J.-I., Ganesan, S., Luo, S.X., Wu, Y.-W., Park, E., Huang, E.J., Chen, L., and Ding, J.B. (2015). Aldehyde dehydrogenase 1a1 mediates a GABA synthesis pathway in midbrain dopaminergic neurons. Science 350, 102–106.

Kumar, R., Corbett, M.A., van Bon, B.W.M., Woenig, J.A., Weir, L., Douglas, E., Friend, K.L., Gardner, A., Shaw, M., Jolly, L.A., et al. (2015). THOC2 Mutations Implicate mRNA-Export Pathway in X-Linked Intellectual Disability. Am. J. Hum. Genet. 97, 302–310.

Kutsarova, E., Munz, M., and Ruthazer, E.S. (2017). Rules for Shaping Neural Connections in the Developing Brain. Front. Neural Circuits 10.

Lakhina, V., Arey, R.N., Kaletsky, R., Kauffman, A., Stein, G., Keyes, W., Xu, D., and Murphy, C.T. (2015). Genome-wide Functional Analysis of CREB/Long-Term Memory-Dependent Transcription Reveals Distinct Basal and Memory Gene Expression Programs. Neuron 85, 330–345.

Laßek, M., Weingarten, J., and Volknandt, W. (2015). The synaptic proteome. Cell Tissue Res. 359, 255–265.

Lek, M., Karczewski, K.J., Minikel, E.V., Samocha, K.E., Banks, E., Fennell, T., O’Donnell-Luria, A.H., Ware, J.S., Hill, A.J., Cummings, B.B., et al. (2016). Analysis of protein-coding genetic variation in 60,706 humans. Nature 536, 285–291.

Lenzken, S.C., Achsel, T., Carrì, M.T., and Barabino, S.M.L. (2014). Neuronal RNA-binding proteins in health and disease. Wiley Interdiscip. Rev. RNA 5, 565–576.

Lill, C.M., Abel, O., Bertram, L., and Al-Chalabi, A. (2011). Keeping up with genetic discoveries in amyotrophic lateral sclerosis: The ALSoD and ALSGene databases. Amyotroph. Lateral Scler. 12, 238–249.

Lill, C.M., Roehr, J.T., McQueen, M.B., Kavvoura, F.K., Bagade, S., Schjeide, B.-M.M., Schjeide, L.M., Meissner, E., Zauft, U., Allen, N.C., et al. (2012). Comprehensive Research Synopsis and Systematic Meta-Analyses in Parkinson’s Disease Genetics: The PDGene Database. PLOS Genet. 8, e1002548.

Liu, R., Hannenhalli, S., and Bucan, M. (2009). Motifs and cis-regulatory modules mediating the expression of genes co-expressed in presynaptic neurons. Genome Biol. 10, R72.

Luna, R., Rondón, A.G., and Aguilera, A. (2012). New clues to understand the role of THO and other functionally related factors in mRNP biogenesis. Biochim. Biophys. Acta BBA -Gene Regul. Mech. 1819, 514–520.

Mancini, A., Niemann-Seyde, S.C., Pankow, R., Omar, E.B., Klebba-Färber, S., Koch, A., Jaworska, E., Elaine, S., Gruber, A.D., Whetton, A.D., et al. (2010). THOC5/FMIP, an mRNA export TREX complex protein, is essential for hematopoietic primitive cell survival in vivo. BMC Biol. 8.

Masuda, S., Das, R., Cheng, H., Hurt, E., Dorman, N., and Reed, R. (2005). Recruitment of the human TREX complex to mRNA during splicing. Genes Dev. 19, 1512–1517.

Misgeld, T., Burgess, R.W., Lewis, R.M., Cunningham, J.M., Lichtman, J.W., and Sanes, J.R. (2002). Roles of Neurotransmitter in Synapse Formation. Neuron 36, 635–648.

Nousiainen, H.O., Kestilä, M., Pakkasjärvi, N., Honkala, H., Kuure, S., Tallila, J., Vuopala, K., Ignatius, J., Herva, R., and Peltonen, L. (2008). Mutations in mRNA export mediator GLE1 result in a fetal motoneuron disease. Nat. Genet. 40, 155–157.

Okuno, H. (2011). Regulation and function of immediate-early genes in the brain: Beyond neuronal activity markers. Neurosci. Res. 69, 175–186.

Patel, M.R., Lehrman, E.K., Poon, V.Y., Crump, J.G., Zhen, M., Bargmann, C.I., and Shen, K. (2006). Hierarchical assembly of presynaptic components in defined C. elegans synapses. Nat. Neurosci. 9, 1488–1498.

Peña, Á., Gewartowski, K., Mroczek, S., Cuéllar, J., Szykowska, A., Prokop, A., Czarnocki-Cieciura, M., Piwowarski, J., Tous, C., Aguilera, A., et al. (2012). Architecture and nucleic acids recognition mechanism of the THO complex, an mRNP assembly factor. EMBO J. 31, 1605–1616.

Piggott, B.J., Liu, J., Feng, Z., Wescott, S.A., and Xu, X.Z.S. (2011). The Neural Circuits and Synaptic Mechanisms Underlying Motor Initiation in C. elegans. Cell 147, 922–933.

Pitzonka, L., Ullas, S., Chinnam, M., Povinelli, B.J., Fisher, D.T., Golding, M., Appenheimer, M.M., Nemeth, M.J., Evans, S., and Goodrich, D.W. (2014). The Thoc1 Encoded Ribonucleoprotein Is Required for Myeloid Progenitor Cell Homeostasis in the Adult Mouse. PLoS ONE 9, e97628.

Pokala, N., Liu, Q., Gordus, A., and Bargmann, C.I. (2014). Inducible and titratable silencing of Caenorhabditis elegans neurons in vivo with histamine-gated chloride channels. Proc. Natl. Acad. Sci. 111, 2770–2775.

Prakash, N., and Wurst, W. (2006). Development of dopaminergic neurons in the mammalian brain. Cell. Mol. Life Sci. CMLS 63, 187–206.

Raj, B., and Blencowe, B.J. (2015). Alternative Splicing in the Mammalian Nervous System: Recent Insights into Mechanisms and Functional Roles. Neuron 87, 14–27.

Rangaraju, V., Dieck, S. tom, and Schuman, E.M. (2017). Local translation in neuronal compartments: how local is local? EMBO Rep. e201744045.

Rehwinkel, J., Herold, A., Gari, K., Köcher, T., Rode, M., Ciccarelli, F.L., Wilm, M., and Izaurralde, E. (2004). Genome-wide analysis of mRNAs regulated by the THO complex in Drosophila melanogaster. Nat. Struct. Mol. Biol. 11, 558–566.

Riccomagno, M.M., and Kolodkin, A.L. (2015). Sculpting Neural Circuits by Axon and Dendrite Pruning. Annu. Rev. Cell Dev. Biol. 31, 779–805.

Sakamoto, K., Karelina, K., and Obrietan, K. (2011). CREB: a multifaceted regulator of neuronal plasticity and protection. J. Neurochem. 116, 1–9.

Saran, S., Tran, D.D.H., Ewald, F., Koch, A., Hoffmann, A., Koch, M., Nashan, B., and Tamura, T. (2016). Depletion of three combined THOC5 mRNA export protein target genes synergistically induces human hepatocellular carcinoma cell death. Oncogene 35, 3872–3879.

Saran, Shashank, Tran, Doan DH, Klebba-Färber, Sabine, Moran-Losada, Patricia, Wiehlmann, Lutz, Koch, Alexandra, Chopra, Himpriya, Pabst, Oliver, Hoffmann, Andrea, Kopfleisch, Robert, et al. (2013). THOC5, a member of the mRNA export complex, contributes to processing of a subset of wingless/ integrated (Wnt) target mRNAs and integrity of the gut epithelial barrier. BMC Cell Biol. 14.

Sawin, E.R., Ranganathan, R., and Horvitz, H.R. (2000). C. elegans Locomotory Rate Is Modulated by the Environment through a Dopaminergic Pathway and by Experience through a Serotonergic Pathway. Neuron 26, 619–631.

Schwartz, M.L., and Jorgensen, E.M. (2016). SapTrap, a Toolkit for High-Throughput CRISPR/Cas9 Gene Modification in Caenorhabditis elegans. Genetics 202, 1277–1288.

Sepuri, N.B.V., Tammineni, P., Mohammed, F., and Paripati, A. (2016). Nuclear Transcription Factors in the Mitochondria: A New Paradigm in Fine-Tuning Mitochondrial Metabolism. (Springer Berlin Heidelberg), pp. 1–18.

Seytanoglu, A., Alsomali, N.I., Valori, C.F., McGown, A., Kim, H.R., Ning, K., Ramesh, T., Sharrack, B., Wood, J.D., and Azzouz, M. (2016). Deficiency in the mRNA export mediator Gle1 impairs Schwann cell development in the zebrafish embryo. Neuroscience 322, 287–297.

Sørensen, B.B., Ehrnsberger, H.F., Esposito, S., Pfab, A., Bruckmann, A., Hauptmann, J., Meister, G., Merkl, R., Schubert, T., Längst, G., et al. (2017). The Arabidopsis THO/TREX component TEX1 functionally interacts with MOS11 and modulates mRNA export and alternative splicing events. Plant Mol. Biol. 93, 283–298.

Spitzer, N.C. (2012). Activity-dependent neurotransmitter respecification. Nat. Rev. Neurosci. 13, 94–106.

Stefanakis, N., Carrera, I., and Hobert, O. (2015). Regulatory Logic of Pan-Neuronal Gene Expression in C. elegans. Neuron 87, 733–750.

Sträßer, K., Masuda, S., Mason, P., Pfannstiel, J., Oppizzi, M., Rodriguez-Navarro, S., Rondón, A.G., Aguilera, A., Struhl, K., Reed, R., et al. (2002). TREX is a conserved complex coupling transcription with messenger RNA export. Nature 417, 304–308.

Tran, D.D., Saran, S., Dittrich-Breiholz, O., Williamson, A.J., Klebba-Färber, S., Koch, A., Kracht, M., Whetton, A.D., and Tamura, T. (2013). Transcriptional regulation of immediate-early gene response by THOC5, a member of mRNA export complex, contributes to the M-CSF-induced macrophage differentiation. Cell Death Dis. 4, e879.

Tran, D.D.H., Saran, S., Williamson, A.J.K., Pierce, A., Dittrich-Breiholz, O., Wiehlmann, L., Koch, A., Whetton, A.D., and Tamura, T. (2014a). THOC5 controls 31end-processing of immediate early genes via interaction with polyadenylation specific factor 100 (CPSF100). Nucleic Acids Res. 42, 12249–12260.

Tran, D.D.H., Koch, A., and Tamura, T. (2014b). THOC5, a member of the mRNA export complex: a novel link between mRNA export machinery and signal transduction pathways in cell proliferation and differentiation. Cell Commun. Signal.

Tritsch, N.X., Ding, J.B., and Sabatini, B.L. (2012). Dopaminergic neurons inhibit striatal output through non-canonical release of GABA. Nature 490, 262–266.

Tsoi, H., Lau, C.K., Lau, K.F., and Chan, H.Y.E. (2011). Perturbation of U2AF65/NXF1-mediated RNA nuclear export enhances RNA toxicity in polyQ diseases. Hum. Mol. Genet. 20, 3787–3797.

Van Vactor, D., and Sigrist, S.J. (2017). Presynaptic morphogenesis, active zone organization and structural plasticity in Drosophila. Curr. Opin. Neurobiol. 43, 119–129.

Voynov, V., Verstrepen, K.J., Jansen, A., Runner, V.M., Buratowski, S., and Fink, G.R. (2006). Genes with internal repeats require the THO complex for transcription. Proc. Natl. Acad. Sci. U. S. A. 103, 14423–14428.

Wang, L., Miao, Y.-L., Zheng, X., Lackford, B., Zhou, B., Han, L., Yao, C., Ward, J.M., Burkholder, A., Lipchina, I., et al. (2013). The THO Complex Regulates Pluripotency Gene mRNA Export and Controls Embryonic Stem Cell Self-Renewal and Somatic Cell Reprogramming. Cell Stem Cell 13, 676–690.

Wang, X., Chang, Y., Li, Y., Zhang, X., and Goodrich, D.W. (2006). Thoc1/Hpr1/p84 is essential for early embryonic development in the mouse. Mol. Cell. Biol. 26, 4362–4367.

Wang, X., Chinnam, M., Wang, J., Wang, Y., Zhang, X., Marcon, E., Moens, P., and Goodrich, D.W. (2009). Thoc1 Deficiency Compromises Gene Expression Necessary for Normal Testis Development in the Mouse. Mol. Cell. Biol. 29, 2794–2803.

White, J.G., Southgate, E., Thomson, J.N., and Brenner, S. (1986). The Structure of the Nervous System of the Nematode Caenorhabditis elegans. Philos. Trans. R. Soc. B Biol. Sci. 314, 1–340.

Woerner, A.C., Frottin, F., Hornburg, D., Feng, L.R., Meissner, F., Patra, M., Tatzelt, J., Mann, M., Winklhofer, K.F., Hartl, F.U., et al. (2016). Cytoplasmic protein aggregates interfere with nucleocytoplasmic transport of protein and RNA. Science 351, 173–176.

Yin, J.C.P., Wallach, J.S., Del Vecchio, M., Wilder, E.L., Zhou, H., Quinn, W.G., and Tully, T. (1994). Induction of a dominant negative CREB transgene specifically blocks long-term memory in Drosophila. Cell 79, 49–58.

Zhang, K., Donnelly, C.J., Haeusler, A.R., Grima, J.C., Machamer, J.B., Steinwald, P., Daley, E.L., Miller, S.J., Cunningham, K.M., Vidensky, S., et al. (2015). The C9orf72 repeat expansion disrupts nucleocytoplasmic transport. Nature 525, 56–61.

Zigmond, M.J., Acheson, A.L., Stachowiak, M.K., and Strickerm, E.M. (1984). Neurochemical Compensation After Nigrostriatal Bundle Injury in an Animal Model of Preclinical Parkinsonism. Arch. Neurol. 41, 856–861.

