## Supplementary figures and images for "A conserved nuclear export complex coordinates transcripts for dopaminergic synaptogenesis and neuronal surviva"

### Supplementary Materials

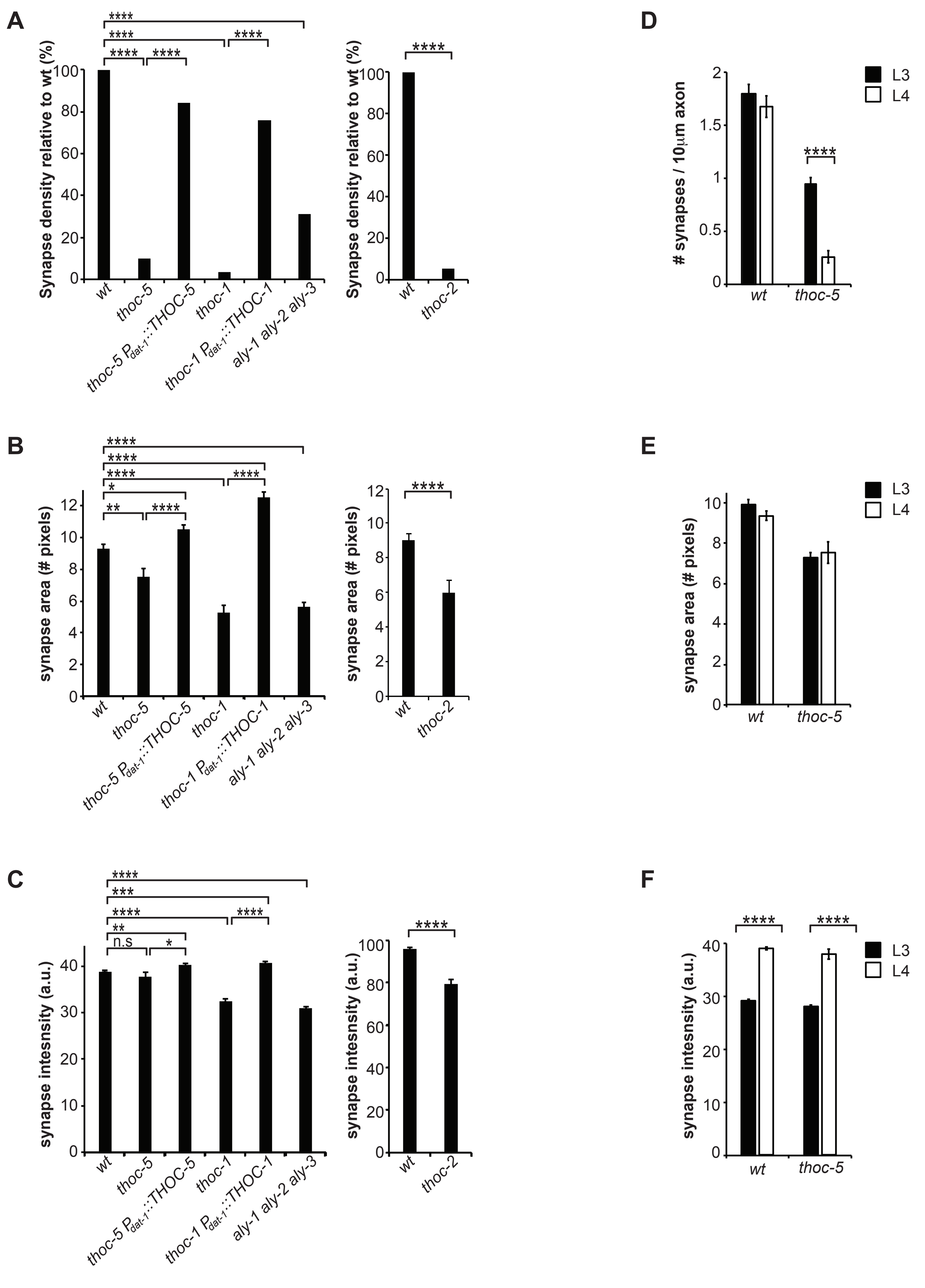

### Supplementary Materials

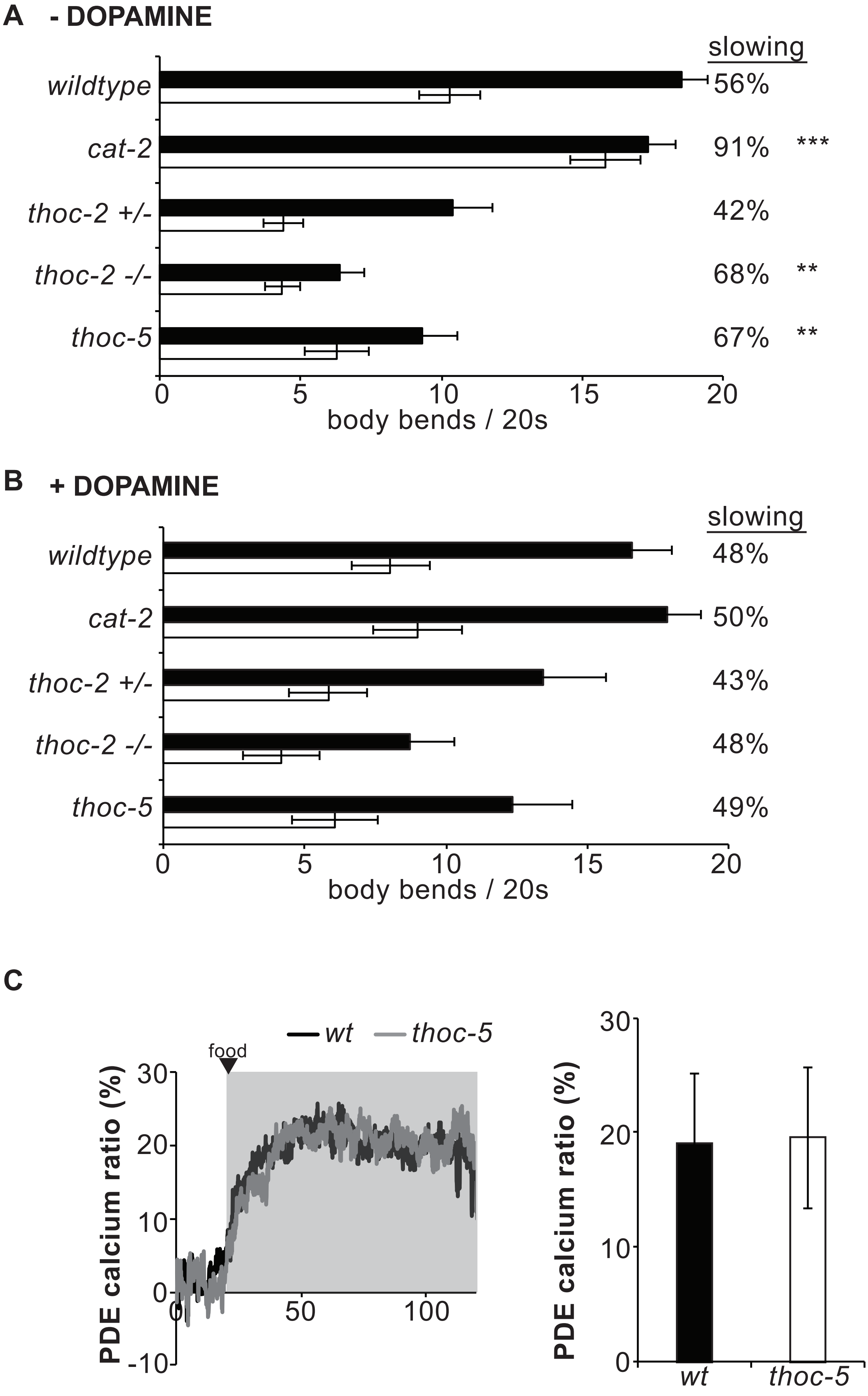

### Supplementary Materials

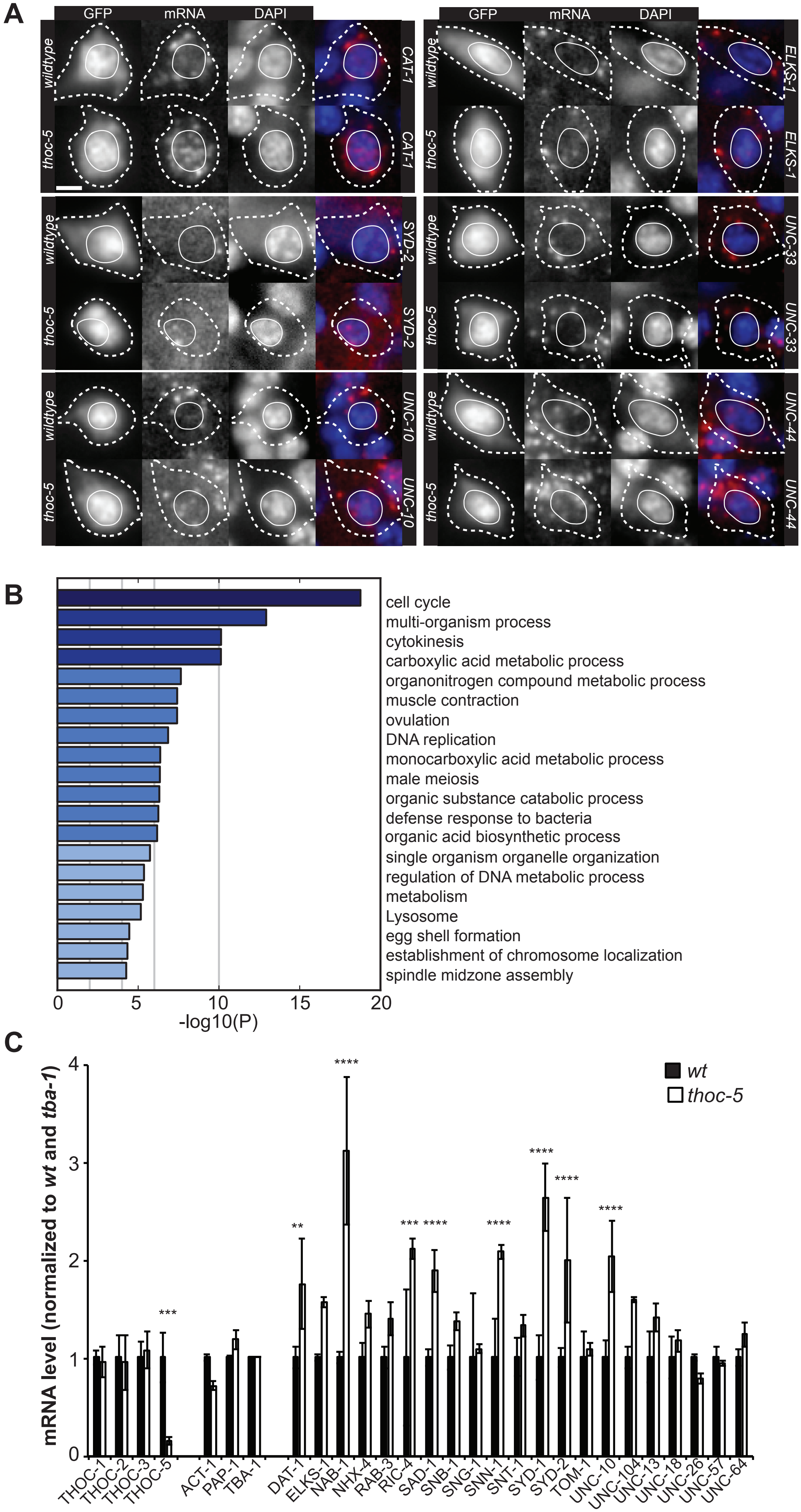

### Supplementary Materials

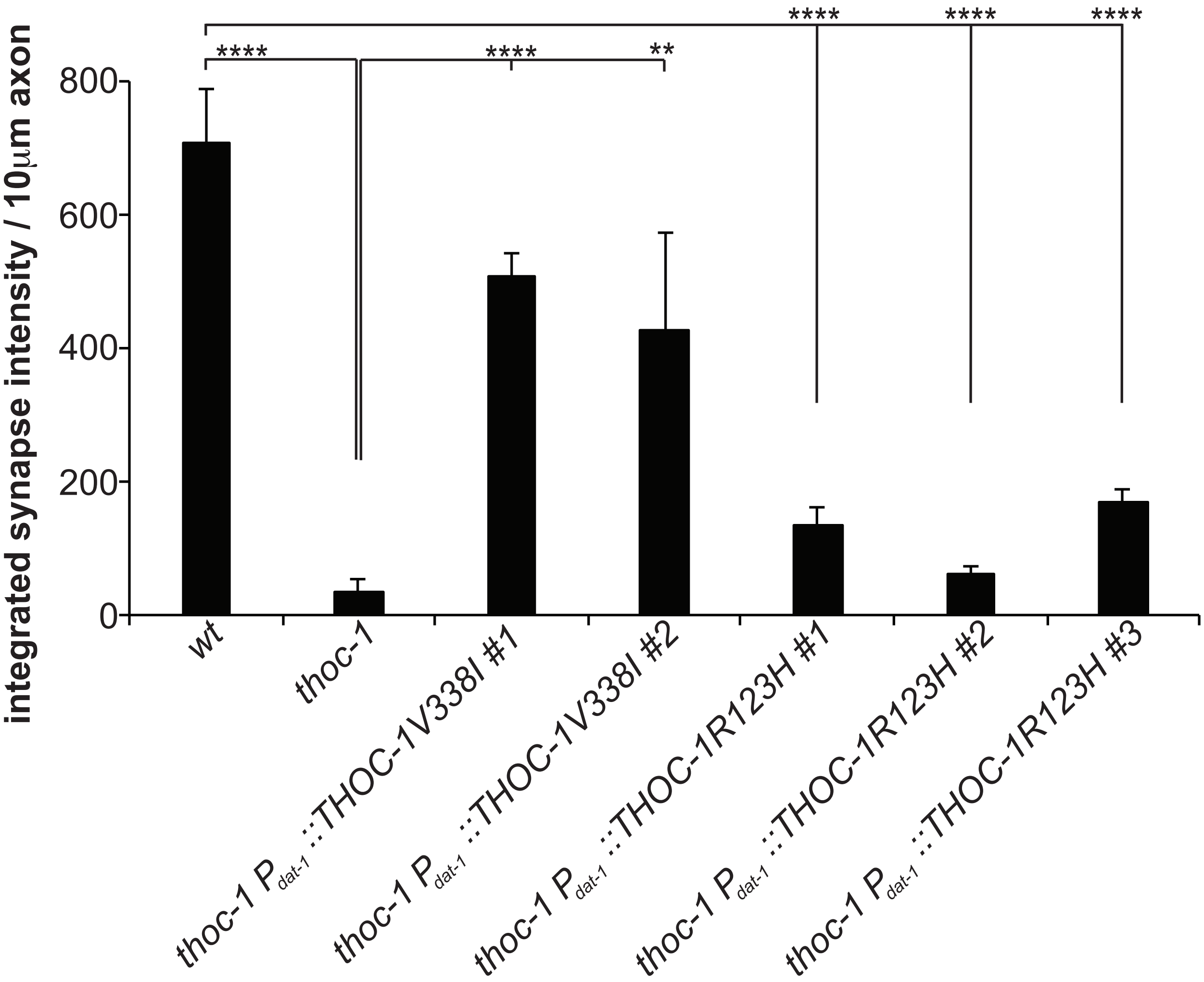

### Supplementary Materials

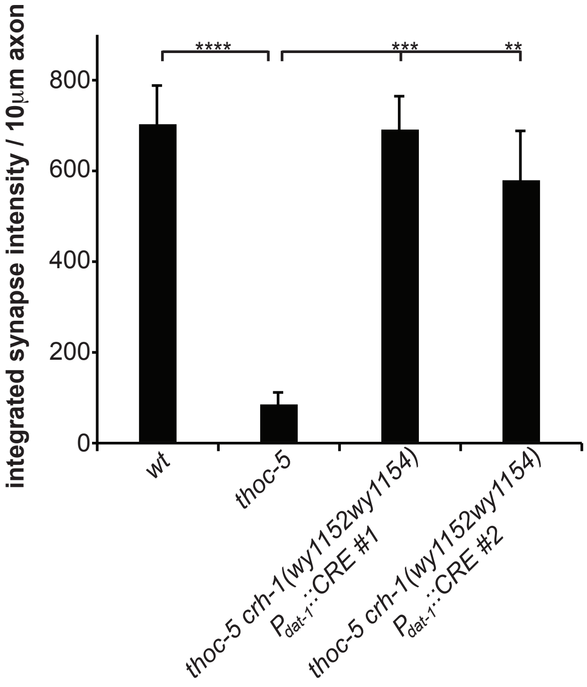

### Supplementary Materials

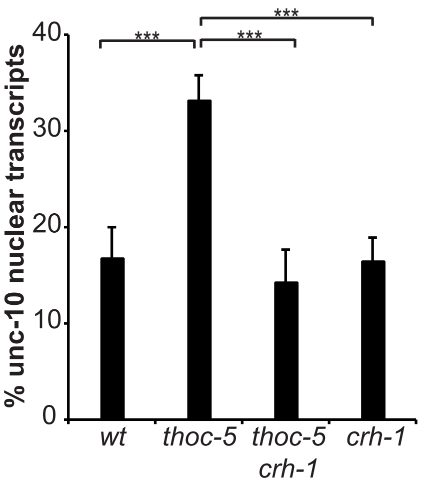

### Supplementary Materials

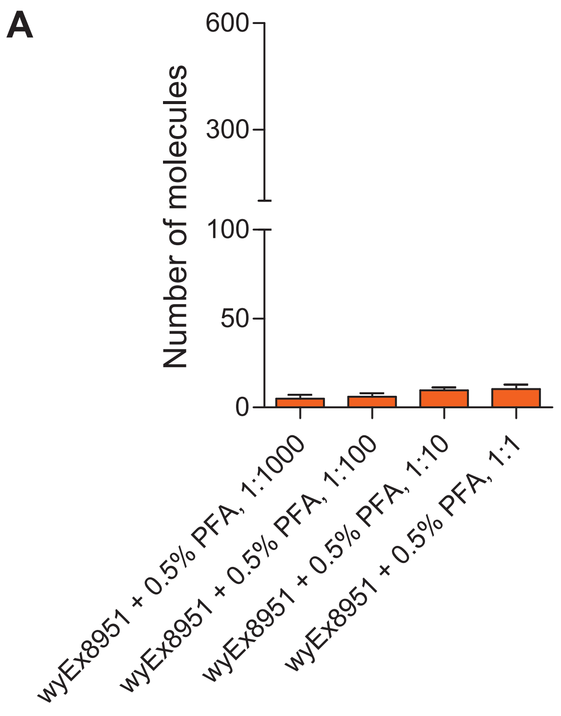

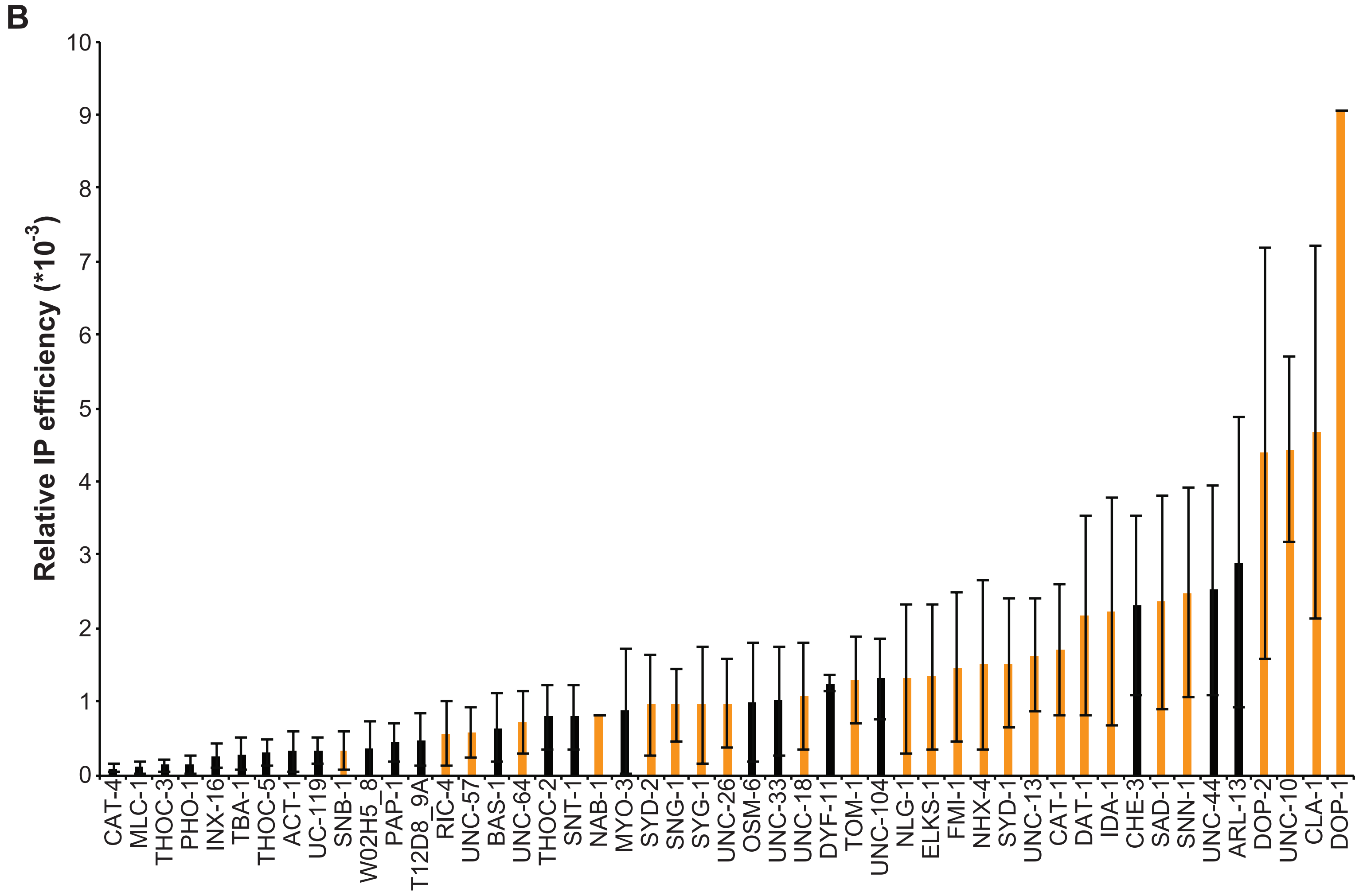
